# Evolutionary origins and diversification of testis-specific short histone H2A variants in mammals

**DOI:** 10.1101/165936

**Authors:** Antoine Molaro, Janet M. Young, Harmit S. Malik

**Affiliations:** Division of Basic Sciences, Fred Hutchinson Cancer Research Center, Seattle, WA 98109; Howard Hughes Medical Institute, Fred Hutchinson Cancer Research Center, Seattle, WA 98109

## Abstract

Eukaryotic genomes must accomplish the tradeoff between compact packaging for genome stability and inheritance, and accessibility for gene expression. They do so using post-translational modifications of four ancient canonical histone proteins (H2A, H2B, H3 and H4), and by deploying histone variants with specialized chromatin functions. While some histone variants are highly conserved across eukaryotes, others carry out lineage-specific functions. Here, we characterize the evolution of male germline-specific “short H2A variants”, which wrap shorter DNA fragments than canonical H2A. In addition to three previously described H2A.B, H2A.L and H2A.P variants, we describe a novel, extremely short H2A histone variant: H2A.Q. We show that *H2A.B, H2A.L, H2A.P* and *H2A.Q* are most closely related to a novel, more canonical mmH2A variant found only in monotremes and marsupials. Using phylogenomics, we trace the origins and early diversification of short histone variants into four distinct clades to the ancestral X chromosome of placental mammals. We show that short H2A variants further diversified by repeated lineage-specific amplifications and losses, including pseudogenization of *H2A.L* in many primates. We also uncover evidence for concerted evolution of *H2A.B* and *H2A.L* genes by gene conversion in many species, involving loci separated by large distances. Finally, we find that short H2As evolve more rapidly than any other histone variant, with evidence that positive selection has acted upon *H2A.P* in primates. Based on their X chromosomal location and pattern of genetic innovation, we speculate that short H2A histone variants are engaged in a form of genetic conflict involving the mammalian sex chromosomes.

## Introduction

Nucleosomes are the basic unit of chromatin in practically all eukaryotes. A typical nucleosome particle wraps 150bp of DNA around an octamer of four histone proteins: H3, H4, H2A and H2B (Kornberg 1974; Kornberg and Thomas 1974; Malik and Henikoff 2003). These four canonical “core” histone proteins have a stereotypical structure characterized by alpha-helices comprising a “histone fold domain” (HFD) (Luger, et al. 1997). During nucleosome assembly, the HFD mediates dimerization of H3 with H4, and that of H2A with H2B, and contains the nucleosome-DNA interface. Nucleosomes thus comprise a central core of a H3-H4 tetramer, flanked by two H2A-H2B dimers. In contrast to their HFD, the N-and C-terminal tails of canonical histones are far less structured and remain solvent-exposed in the nucleosome.

Nucleosomes can prevent other cellular factors, such as transcription factors, from interacting with DNA. While the post-translational modification of histone tails play an important role in regulating these interactions, eukaryotic genomes also establish diverse chromatin states using histone variants that replace canonical histones (Weber and Henikoff 2014; Talbert and Henikoff 2017). Although histone variants are closely related to canonical histones, they differ at several key amino acid residues that dictate chromatin states; many histone variants also contain additional domains essential for their non-canonical functions (Malik and Henikoff 2003; Talbert and Henikoff 2010; Draizen, et al. 2016). Altering histone composition greatly influences nucleosome compaction and regulation of chromatin function. For example, the H2A.Z variant is incorporated at transcriptional start sites of genes to facilitate RNA polymerase II (RNAPII) release, the cenH3 variant triggers the assembly of the kinetochore at centromeric heterochromatin, and the H2A.X variant assists with DNA repair at sites of double-strand breaks (Weber and Henikoff 2014; Talbert and Henikoff 2017).

From an evolutionary standpoint, eukaryotic canonical histones are thought to derive from histones found in some archaea (Sandman and Reeve 2006), and are some of the most conserved proteins across eukaryotes. Canonical histones H3, H4 and H2A have evolved under very strong purifying selection; each class of histones has nearly identical HFDs across all eukaryotes (>80% a.a. identity) (Malik and Henikoff 2003; Draizen, et al. 2016). Although H2B proteins display more divergence, they remain exceptionally well conserved since the origins of eukaryotes (>50% a.a. identity) (Malik and Henikoff 2003; Gonzalez-Romero, et al. 2010). Canonical histone genes are usually present in multiple copies in genomes, arranged in multi-gene clusters, but the genomic organization and copy number varies between species (Marzluff, et al. 2008).

Unlike canonical histones, histone variants are encoded by “stand-alone” single-copy genes and originated during the radiation of eukaryotes. Some variants, such as H2A.Z and cenH3, diverged from canonical histones early in eukaryotic evolution. Other variants, such as H2A.X, might have arisen independently in several lineages, whereas variants such as macroH2A are present only in some lineages (Malik and Henikoff 2003; Talbert and Henikoff 2010; Rivera-Casas, et al. 2016). Despite these diverse origins, most histone variants evolve under strong purifying selection (Piontkivska, et al. 2002; Rooney, et al. 2002; Talbert, et al. 2012). Only one surprising exception has been well characterized: the centromere-specific H3 variant, cenH3. Phylogenetic studies have shown that closely related species of plants and animals have surprisingly divergent cenH3 sequences that are subject to recurrent diversifying, or positive, selection (Malik and Henikoff 2001; Talbert, et al. 2002). This finding of rapid evolution is a hallmark of evolutionary arms races between interacting genetic elements with opposite evolutionary interests (Henikoff and Malik 2002; McLaughlin and Malik 2017). In this case, the rapid evolution of cenH3 variants is hypothesized to be a result of “centromere drive”, in which allelic centromeres adapt to compete for oocyte inclusion after female meiosis, and drive counter-adaptation by centromere proteins such as cenH3 to suppress the driving elements (Henikoff, et al. 2001; Fishman and Saunders 2008; Malik and Henikoff 2009; Chmatal, et al. 2014).

During mammalian male germ cell development, unique sets of histone variants populate the chromatin of meiotic and post-meiotic germ cells (Boussouar, et al. 2008). This reprogramming culminates with the nearly complete replacement of histones by protamines, leaving only a few nucleosomes behind (Oliva and Dixon 1991; Hammoud, et al. 2009). Among the histone variants that are expressed during this developmental window, one family known as “short H2A variants” remains poorly characterized (Govin, et al. 2007; Ferguson, et al. 2009). Although short H2As display unusual sequence properties, their evolutionary origins and trajectories are still not fully understood (Eirin-Lopez, et al. 2008; Ishibashi, et al. 2010).

Three short H2A variants, H2A.B, H2A.L and H2A.P, have been described so far; each exhibits less than 50% amino acid identity with canonical H2A (Shaytan, et al. 2015; Draizen, et al. 2016). Both *H2A.B* and *H2A.L* are found in multiple copies in the mouse and human genomes (Govin, et al. 2007; Ishibashi, et al. 2010). Although one study has also detected expression of H2A.B in mouse embryonic stem cells (Chen, et al. 2014), short histone H2A variant expression has been primarily observed in the male germline. During mouse spermatogenesis, H2A.B is first detected in spermatocyte nuclei at the onset of meiosis (Soboleva, et al. 2012). At the end of meiosis, H2A.B staining disappears whereas H2A.L accumulates in spermatid nuclei until the end of spermatogenesis (Govin, et al. 2007). Importantly, some H2A.L remains in mature sperm chromatin as part of the nucleosome complement that is refractory to protamine exchange in mouse (Govin, et al. 2007; Baker, et al. 2008; Ishibashi, et al. 2010). However, following fertilization, mouse H2A.L staining is rapidly lost from the paternal pronucleus (Wu, et al. 2008). In contrast, human sperm lacks H2A.L protein, and instead retains H2A.B (Baker, et al. 2007). Although *H2A.P* RNA is clearly enriched in mouse testes, H2A.P protein has not yet been detected (Govin, et al. 2007; El Kennani, et al. 2017).

Relative to canonical H2A, the three short H2A variants have truncated C-termini, shortening the “docking domain” that mediates the interaction with H3-H4 dimers, and eliminating the lysine residue (K119), whose ubiquitination normally regulates H2A turnover (Figure 1) (Bonisch and Hake 2012; Buschbeck and Hake 2017). Most of the N-terminal tail’s lysines (K), which are typically subject to post-translational modifications in canonical histones, have been replaced by arginine (R). Finally, short H2As have alterations in their acidic patch, which normally stabilizes nucleosome-nucleosome interactions during chromatin compaction in other H2As (Luger, et al. 1997; Bonisch and Hake 2012; Soboleva, et al. 2012; Shaytan, et al. 2015; Buschbeck and Hake 2017). Based on this feature, H2A.B was previously also referred to as H2A.Lap1 for “lack of acidic patch” (Soboleva, et al. 2012). Consequently, nucleosomes containing short H2A variants wrap shorter stretches of DNA (~120-130bp) than canonical H2A-containing nucleosomes (~150bp) and form more loosely-packed, labile chromatin *in vitro* (Angelov, et al. 2004; Bao, et al. 2004; Gautier, et al. 2004; Doyen, et al. 2006; Syed, et al. 2009; Soboleva, et al. 2012; Arimura, et al. 2013) and *in vivo* (Govin, et al. 2007; Barral, et al. 2017; Soboleva, et al. 2017).

**Figure 1:**
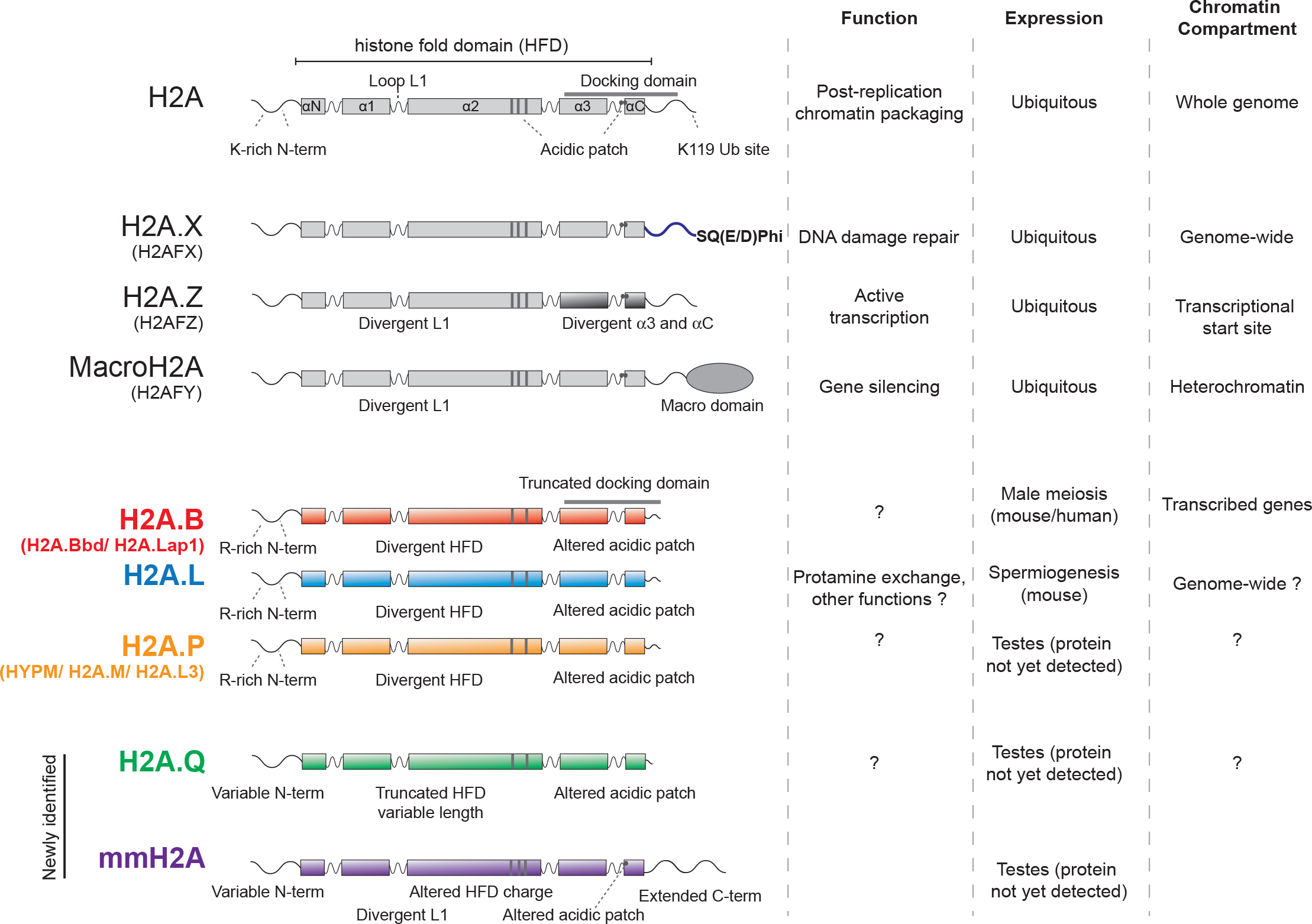
Features of H2A protein families. Alternative names for each family are shown in parentheses. Alpha helices forming the histone fold domain are numbered. The acidic patch is schematized with grey bars and dots in alpha2 and alphaC. The known function, expression and chromatin compartment of these histones are shown on the right. Short H2A variants are colored. “SQ(E/D)phi” denotes the phosphorylation domain of H2A.X where “phi” stands for any hydrophobic residue. Note that mmH2A and H2A.Q are novel variants that are described for the first time in this study.

These structurally divergent features greatly influence the chromatin attributes and functions associated with short H2As. H2A.B nucleosomes are believed to create open chromatin suitable for transcription (Chadwick and Willard 2001; Soboleva, et al. 2012; Tolstorukov, et al. 2012; Soboleva, et al. 2017). Indeed, studies using ectopic expression in mouse and human cell lines have shown that H2A.B is incorporated in nucleosomes and localizes to sites of open chromatin, DNA synthesis and splicing (Chadwick and Willard 2001; Tolstorukov, et al. 2012; Sansoni, et al. 2014). Recent *in vivo* studies, using mice lacking a specific *H2A.L* gene, demonstrate that it is essential for fertility and indicate that it is involved in initiating the replacement of histones with protamines (Barral, et al. 2017). However, the functions of other short H2As are largely unknown.

Several studies have examined which chromatin compartments contain short H2A variants. H2A.B was originally named H2A.Bbd, for “Barr body deficient” (Chadwick and Willard 2001) because it is excluded from the inactive X chromosome (the “Barr body”) when ectopically expressed in female cell lines. ChIP-seq experiments using mouse testis chromatin revealed H2A.B accumulation at the transcriptional start site and body of active genes (Soboleva, et al. 2012). In addition, H2A.B was shown to directly interact with the transcriptional machinery as well as RNA (Soboleva, et al. 2017). In mouse ES cells, H2A.B protein seems to be found in the chromatin of imprinted genes (Chen, et al. 2014). Although initial immunofluorescence studies suggested that mouse H2A.L proteins reside in pericentromeric regions of post-meiotic spermatid nuclei, recent mapping experiments found a much broader genome-wide distribution (Govin, et al. 2007; Barral, et al. 2017).

The lack of a comprehensive evolutionary analysis of the three known short H2A variants has created confusion both in nomenclature and the interpretation of functional studies. For instance, *H2A.P* was initially believed to be an allele of *H2A.L* (Govin, et al. 2007), but has more recently been confirmed as a distinct histone variant (Marino-Ramirez, et al. 2006; Draizen, et al. 2016; El Kennani, et al. 2017). Lack of a phylogenetic framework has also blurred the relationships between short histone variants, particularly between apparent mouse and human orthologs. In order to address these shortcomings, we performed detailed phylogenomic analyses of the short histone variants. Analyses based on our phylogenomic framework allow us to discover two previously undescribed H2A histone variants: mmH2A variants present only in monotremes and marsupials, and H2A.Q variants present in some eutherian mammals. We show that a single precursor to the short H2A variants likely arose in the common ancestor of all eutherian mammals following their divergence from marsupials. This precursor gene duplicated and diverged into the four evolutionarily distinct clades we see today: *H2A.B, H2A.L, H2A.P* and *H2A.Q*. Genes encoding all four variants resided on the X chromosome of the common ancestor of eutherian mammals; *H2A.B* and *H2A.L* each spawned additional ancestral X chromosome linked (X-linked) duplicates. As eutherian mammal lineages continued to diverge, all four clades of short H2A variant genes experienced further diversification by repeated gains and losses while also undergoing lineage-specific concerted evolution between duplicated copies in some cases. As a result, the repertoires of short histone H2A variants vary extensively among eutherian mammals. We show that all four short H2As are subject to accelerated rates of protein evolution relative to both their canonical counterparts and to other known H2A variants. Finally, we find that *H2A.P*, and perhaps *H2A.B*, has evolved under positive selection in simian primates, representing the second class of histone variants to bear such a signature of genetic innovation (after *cenH3*). Taken together, our analyses reveal the origins and evolutionary trajectories of short H2A variants and suggest that these histone variants might be engaged in a novel form of genetic conflict involving mammalian sex chromosomes in the male germline.

## Results

### Age and orthology of short histone H2A variants in eutherian mammals

Previous phylogenetic analyses of short H2A histone variants have mostly focused on *H2A.B* genes, finding that *H2A.B* genes are restricted to mammals and appear to be diverging faster than both canonical histones and other H2A variants (Malik and Henikoff 2003; Eirin-Lopez, et al. 2008; Ishibashi, et al. 2010; Talbert, et al. 2012). Additional studies revealed that mouse *H2A.L* genes experienced lineage-specific amplifications, hinting at the possibility that different mammals might bear different repertoires of short histone variants (Ferguson, et al. 2009). We sought a comparative phylogenetic framework by which we could resolve the initial uncertainty of orthology assignments based on similarity (El Kennani, et al. 2017), as well as elucidate the age and evolutionary dynamics of short H2A variants to gain insights into their possible functions. We scanned genome sequences from representative mammals (and chicken as an outgroup) using BLASTn and tBLASTn searches with human H2A.B, human H2A.P and rat H2A.L sequences as queries (Supp. Table 1). We also examined the genomic location and neighboring genes of each candidate short H2A variant in order to use shared synteny to identify genes that had been present at each genomic location in ancestral mammalian species.

Based on shared syntenic locations and sequence similarity, we were able to unambiguously assign most short H2A variants to be *H2A.B, H2A.P* or *H2A.L* (Figure 2). However, our survey also revealed a single gene in many mammalian genomes (including human) that appeared to be distinct from any of the previously described short histone H2A variants. Although we initially classified it as a divergent H2A.L gene, it became apparent that this gene was quite distinct from all the *bona fide* H2A.L paralogs and maintained as an open reading frame in many but not all eutherian mammals, suggesting it was ancient. It also appeared to encode the shortest H2A variant, consisting almost entirely of just the histone-fold domain (HFD). Based on its conservation in multiple, distinct lineages of eutherian mammals, and the distinctive features of the encoded proteins (see next sections), we tentatively named this novel clade of short histone variants as *H2A.Q*. The putative *H2A.B*, *H2A.L, H2A.P* and *H2A.Q* sequences all have intronless open reading frames (like canonical *H2A*). We were also able to identify pseudogenes that were present in orthologous genomic locations but interrupted by early stop codons or frameshifts. Based on our analysis, we deduce that around 100 million years ago, the common ancestor of eutherian mammals encoded at least eight short H2A variant genes in six distinct X-linked loci, including three *H2A.B* genes, three *H2A.L* genes, a single *H2A.P* gene, and a single *H2A.Q* gene. For clarity, we numbered these ancestral loci relative to their position on each arm of the X chromosome from telomere to centromere. *H2A.B.1.1* and *H2A.B.1.2* are so named since they seem to have arisen from an ancestral duplication within the X chromosome; we were unable to faithfully assign synteny of these two genes in many genomes with draft quality assemblies.

**Figure 2:**
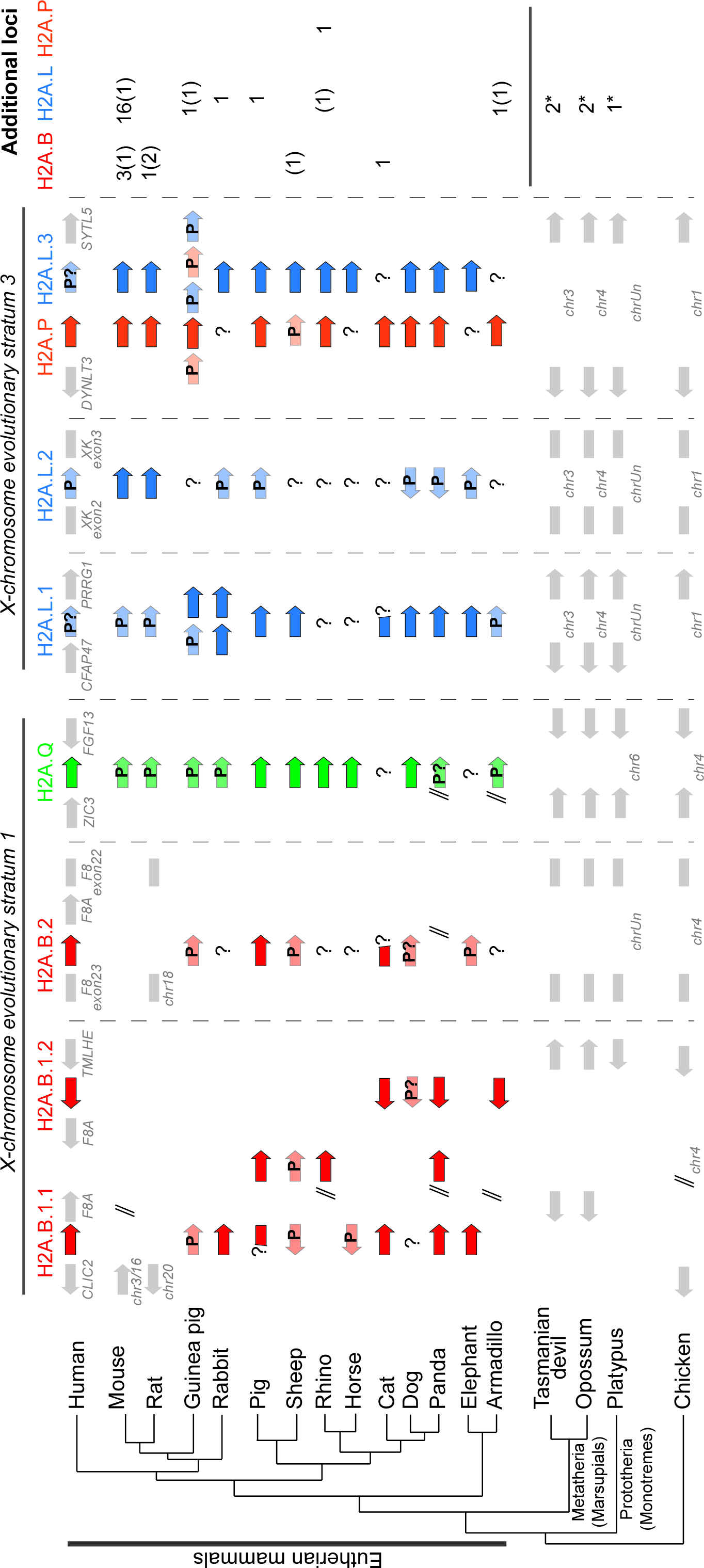
Mammalian short H2A repertoires. We show the eight short H2A loci that existed in the common eutherian ancestor with colored arrows and additional lineage-restricted loci are enumerated on the right (with pseudogenes in parentheses). “*” denotes marsupial and monotreme mmH2A loci. In those genome assemblies where sequences have been mapped to chromosomes, the eight loci containing short H2A variant genes are all on the X chromosome, unless indicated. A species tree of representative mammals is rooted using chicken as an outgroup. Flanking genes (grey arrows) define the syntenic locations of three ancestral *H2A.B* genes (*H2A.B.1.1, H2A.B.1.2* and *H2A.B.2*, red arrows), three ancestral *H2A.L* genes (*H2A.L.1, H2A.L.2* and *H2A.L.3*, blue arrows), *H2A.Q* (green arrow) and *H2A.P* (orange arrow). Species-specific rearrangements within synteny blocks are not indicated for clarity. “P” denotes pseudogenes, “P?” denotes genes with unusually long ORFs that are most likely non-functional (e.g., dog *H2A.B* genes and human *H2A.L* genes); double backslashes indicate assembly gaps that interrupt synteny, and “?” indicates that a gene is not found - in most of these cases, we have not attempted to distinguish true gene loss from absence due to assembly gaps. X chromosome evolutionary strata of these regions are also shown at the top.

In contrast to eutherian mammals, we did not identify clear syntenic orthologs of any of the four short histone H2A variants in marsupial or platypus genomes. However, we did find more distantly related marsupial and platypus H2A variants in non-syntenic locations. To avoid ambiguity, we refer to them as mmH2A (for variants specifically found in <u>m</u>onotremes and <u>m</u>arsupials) and describe them in detail later in this study. Furthermore, we recovered no short histone H2A variants in chicken or any other bird genomes that we queried (zebra finch and turkey). Our analysis allows us to infer that both the origin of short histone variants and their diversification into present-day *H2A.B, H2A.L, H2A.P* and *H2A.Q* occurred in the common ancestor of eutherian mammals after the split from marsupials.

We find that the repertoire of short H2A variants has been greatly shaped by two evolutionary forces. The first of these is recurrent loss. In many species, one or more of the eight ancestral orthologs is now a pseudogene. Indeed, none of the extant mammals that we analyzed still encodes the full original set of short histone variants in syntenic locations, although many have retained evidence of degraded genes at those locations. These pseudogenization events have led to widespread changes in the short H2A histone repertoire of mammals. For instance, all three *H2A.L* copies appear to be pseudogenes in human and some other primates (see below). Furthermore, sheep has a premature stop codon in *H2A.P*, and all *H2A.B* copies are pseudogenes. Finally, all rodents we examined encode a pseudogenized version of *H2A.Q*. Thus, none of the original eutherian short histone H2A gene clades appears to be strictly retained in all mammals, suggesting that neither *H2A.L, H2A.B, H2A.P* nor *H2A.Q* perform universally essential, non-redundant functions in eutherian mammals.

It was not immediately clear whether primate genomes contain any functional *H2A.L* genes. While *H2A.L.2* clearly acquired inactivating mutations early in primate evolution, many primate *H2A.L.1* and *H2A.L.3* genes do not bear any stop codons or frameshifts within the typical *H2A.L* coding region. However, in many primates, the typical H2A.L termination codon has been lost, and the reading frame continues for an additional 18-110 amino acids before the next in-frame stop codon is encountered. In some species, this putative extension of the C-terminus is highly repetitive, and rich in glutamic acid or arginine residues. To determine if any extended *H2A.L* genes are functional, we tested for signals of purifying selection (Supp. Figure 1). We used the codeml algorithm from the PAML package (Yang 1997), to perform maximum likelihood branch tests to ascertain selective pressures among primate *H2A.L* genes. This method compares the rates of synonymous (dS) to non-synonymous (dN) codon substitutions. We found significant evidence (p<0.01) that New World monkey *H2A.L* genes are more likely to be evolving under purifying selection (dN/dS = 0.54) rather than neutrally. In contrast, Old World monkey and hominoid *H2A.L* genes appear to be evolving neutrally (p=0.49, dN/dS=0.80, not statistically distinct from the value expected for a pseudogene of dN/dS=1). Our observations suggest that *H2A.L.1* and *H2A.L.3* are probably functional in New World monkeys, but not in hominoids or Old World monkeys. Our findings support extensive proteomic studies that failed to detect H2A.L protein in human sperm or other tissues, even though H2A.L is readily apparent in mouse sperm (Baker, et al. 2007; Baker, et al. 2008; El Kennani, et al. 2017).

The second force that has dramatically altered short H2A repertoires is lineage-specific duplication and diversification. The most dramatic example of such lineage-specific expansion and diversification has been previously described in mouse (Ferguson, et al. 2009), which encodes a total of 20 *H2A.L* loci, whereas most other mammals encode between zero and five *H2A.L* genes. Although some of these *H2A.L* duplicates may be unique to *Mus musculus*, we found that others were born in or prior to the last common ancestor of all *Mus* species (Supp. Fig. 2).

In genome assemblies where sequences have been assigned to chromosomes, we find that almost all *H2A.B, H2A.L, H2A.P* and *H2A.Q* genes are X-linked. Since the evolution of sex chromosomes involved the progressive suppression of meiotic recombination over large portions of the chromosome arms, one can trace the origin of X-linked genes based on their location along the X chromosome. Short H2A genes are found in “evolutionary strata” known to have resided on the ancestral X chromosome of eutherian mammals (Lahn and Page 1999; Sandstedt and Tucker 2004). This leads us to deduce that *H2A.B, H2A.L, H2A.P* and *H2A.Q* genes likely originated on the X chromosome of eutherian mammals, and have been largely retained on this chromosome. In contrast, the related mmH2A histone variants that we identified in marsupials appear to be autosomal (or on an unmapped scaffold in platypus). The only exceptions to this finding of X-linkage in eutherian mammals are mouse *H2A.L* genes that were previously described as having duplicated/relocated onto the Y chromosome and chromosome 2 (Ferguson, et al. 2009). In the rat genome, it appears that large syntenic rearrangements have moved the loci that historically contained *H2A.B.1.1* and *H2A.B.2* away from the X chromosome; *H2A.B* genes are no longer found in these ancestral, now autosomal loci, and have likely been deleted or pseudogenized beyond recognition. However, the rat X chromosome does still encode an intact *H2A.B* gene. Shared synteny with mouse suggests that this gene (*H2A.B.RatMouse1*, also historically named H2A.B.3 (Soboleva, et al. 2017)) was acquired in the common ancestor of mouse and rat. The loss of autosomal *H2A.B* and retention of X-linked *H2A.B* suggests the intriguing possibility of functional constraint favoring X-linkage. However, we cannot rule out the possibility that X-linkage of short histone variants is largely explained by their evolutionary origins on the X chromosome, rather than by selective retention.

Taken together, our results show that eight short H2A variants were acquired on the X chromosome in the common ancestor of all eutherian mammals following the split with metatherians (marsupials) over 100 million years ago. Following this initial event, our results further show that short H2A variants have experienced a series of lineage-specific events of pseudogenization and duplication. Finally, our analysis indicates that functional H2A short variants have remained largely X-linked during the radiation of eutherian mammals.

### Evolutionary diversification of short H2A variants in mammals

To confirm and extend our findings from the synteny-based analyses, we performed phylogenetic analysis of all H2A canonical and variant proteins (Figure 1). We aligned amino acid sequences of the HFD of canonical and variant H2As (H2A.Z, H2A.X and macroH2A) as well as short histone variants from representative species of mammals and outgroups (chicken and *Xenopus*, Supp. Data 1) and used a maximum likelihood algorithm, PhyML (Guindon and Gascuel 2003; Guindon, et al. 2010) to infer a phylogeny (Figure 3).

**Figure 3:**
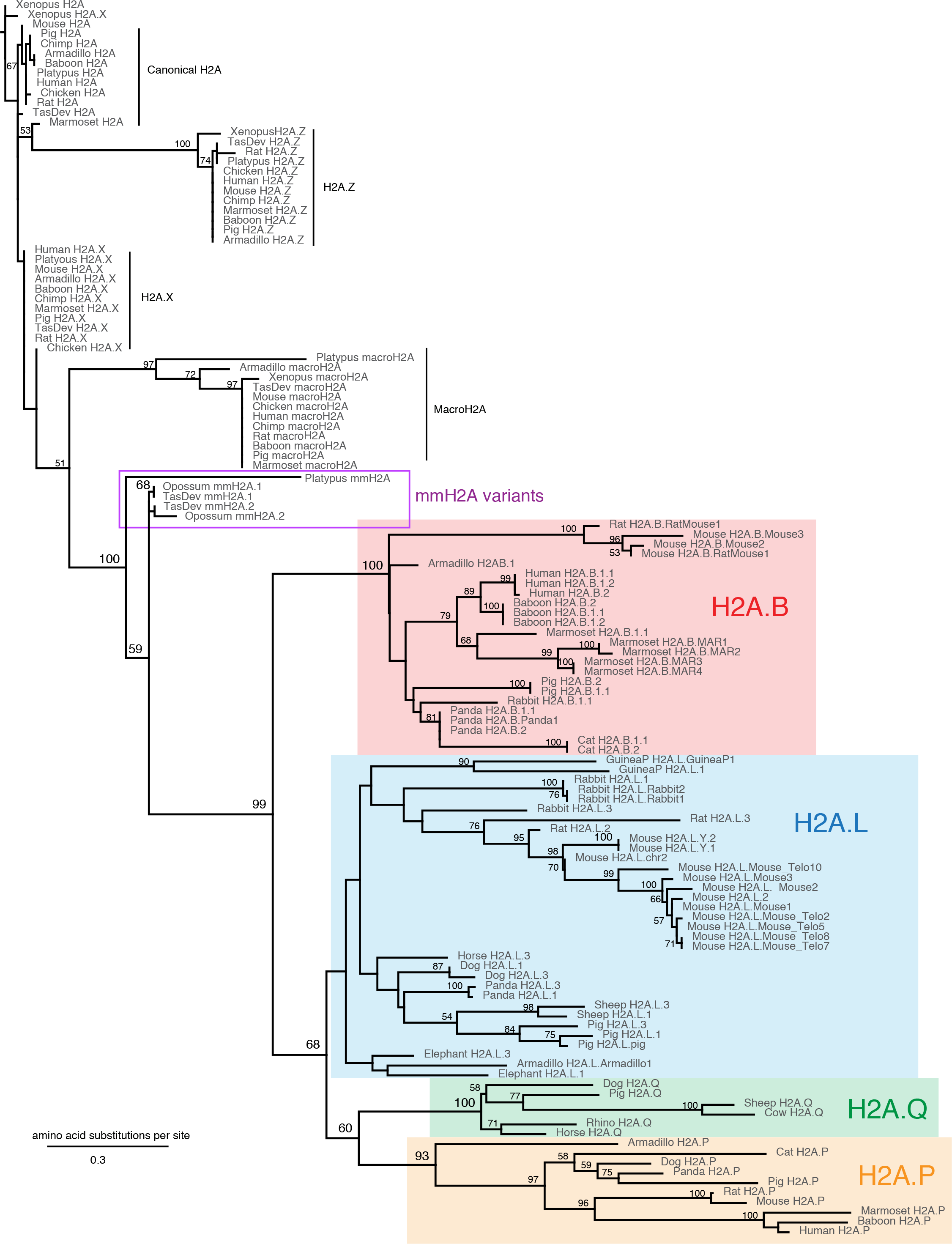
H2A.B, H2A.L, H2A.P, and H2A.Q form distinct clades of H2A variants. Maximum-likelihood protein phylogeny of the histone fold domain of canonical H2A and selected H2A variants. Bootstrap values are shown at all nodes that have >50% bootstrap support. For canonical H2A, H2A.Z, H2A.X and macroH2A, we include sequences from Xenopus, chicken and representative mammalian species. The tree is rooted using Xenopus H2A. Short H2A variant clades are highlighted: H2A.B (red box), H2A.P (orange box), H2A.Q (green box) and H2A.L (blue box). Note that H2A.B, H2A.P and H2A.Q clades are strongly monophyletic whereas the monophyly of H2A.L does not have strong support.

This phylogeny reveals four interesting features of short H2A variants. First, all four short H2A variants (*H2A.B, H2A.L, H2A.P* and *H2A.Q*) form distinct evolutionary clades. This confirms our initial assignment of the *H2A.Q* orthologs as a separate clade, distinct from *H2A.L* and *H2A.P*. *H2A.B, H2A.P* and *H2A.Q* clades all have strong bootstrap values supporting monophyly whereas there is only modest support for *H2A.L*. However, all H2A.L sequences cluster away from other short H2As. Furthermore, the phylogeny suggests that *H2A.P* and *H2A.Q* are more closely related to each other, followed by *H2A.L* and then *H2A.B*. Thus, the phylogenetic analysis confirmed our initial synteny-based assignment of genes as *H2A.B, H2A.L, H2A.P* or *H2A.Q*. The phylogeny also shows that, although they derived from a single common ancestor, *H2A.B, H2A.L, H2A.P* and *H2A.Q* have diverged independently during 100 million years of eutherian mammal evolution.

Second, we found that all eutherian short H2A variants group with the monotreme and marsupial mmH2A variants, to the exclusion of other H2A sequences. The mmH2A variants branch off first and are clearly separated from all eutherian mammal short H2A variants, paralleling the mammalian species tree (Bininda-Emonds, et al. 2007). This suggests that a precursor to the short H2A and mmH2A variants existed in the common ancestor of all mammals, and independently diversified in prototherians (monotremes), metatherians and eutherians.

Third, we see unexpected branching patterns within the *H2A.B* and *H2A.L* clades. Because our synteny analyses showed that the ancestral eutherian mammal had three copies of *H2A.B* and of *H2A.L* in distinct genomic locations, we expected that each genomic copy would diversify on an independent evolutionary trajectory. Under that model, the phylogeny would group *H2A.L.1* genes from different species together, and all *H2A.L.2* genes together, etc. However, we observed a different pattern, whereby all copies of a variant from the same species cluster together in the phylogeny with remarkably high sequence identity. For example, panda *H2A.B.1.1* and *H2A.B.2* share 100% amino acid identity but only 75% identity with their respective syntenic orthologs in cat. These observations suggest the action of sequence homogenization occurring between dispersed histone variant genes within each genome.

Finally, consistent with previous analyses, we found that within the clades representing canonical H2A, H2A.Z, H2A.X and macroH2A, all branch lengths are very short, implying that each is very well-conserved (there is over 90% identity among the species we analyzed). In contrast, branch lengths are much longer within each of the short H2A variant clades, especially in the *H2A.P* and *H2A.Q* clades. This lack of sequence identity either suggests a lower degree of evolutionary constraint, or evolutionary pressure to innovate at the amino acid level.

### Distinguishing structural features of short H2A variant proteins

Our phylogenetic analysis of the four short histone H2A variant types revealed their common evolutionary origins and showed that they acquired changes during early eutherian divergence that clearly distinguish *H2A.B, H2A.L, H2A.P* and *H2A.Q* into four distinct clades. Previous studies have investigated the putative functional and structural consequences of sequence differences in the histone fold domains between canonical H2A and the short H2A variants in mice and humans. To complement these earlier studies, we performed detailed analysis of amino acid conservation in a phylogenetically broad sampling of eutherian mammals to identify protein features common to all short H2A variants but distinct from canonical H2As, as well as features that distinguish short histone H2A variants from one another. Our initial analyses suggested that H2A.Q had many unique features that were not found in any canonical or variant H2A. We therefore initially focused only on the previously described short histone H2A variants *i.e*., H2A.B, H2A.L, and H2A.P, and analyze H2A.Q separately in more detail later.

We created logo plots for canonical H2A, H2A.B, H2A.L, and H2A.P (Figure 4A). For each of these H2A types, we aligned protein sequences from each of eight diverse eutherian mammals, selected so that we could use the same species set for each variant (Methods, Supp. Data 2). Because they reflect the same evolutionary timespan, these logo plots allow us to compare selective constraints between variant types. We aligned these plots to each other where possible, and used the nucleosome crystal structure (PDB:1AOI) (Luger, et al. 1997) to annotate the five alpha-helices and connecting loops of the HFD as well as DNA-proximal and H2B-contacting residues.

**Figure 4:**
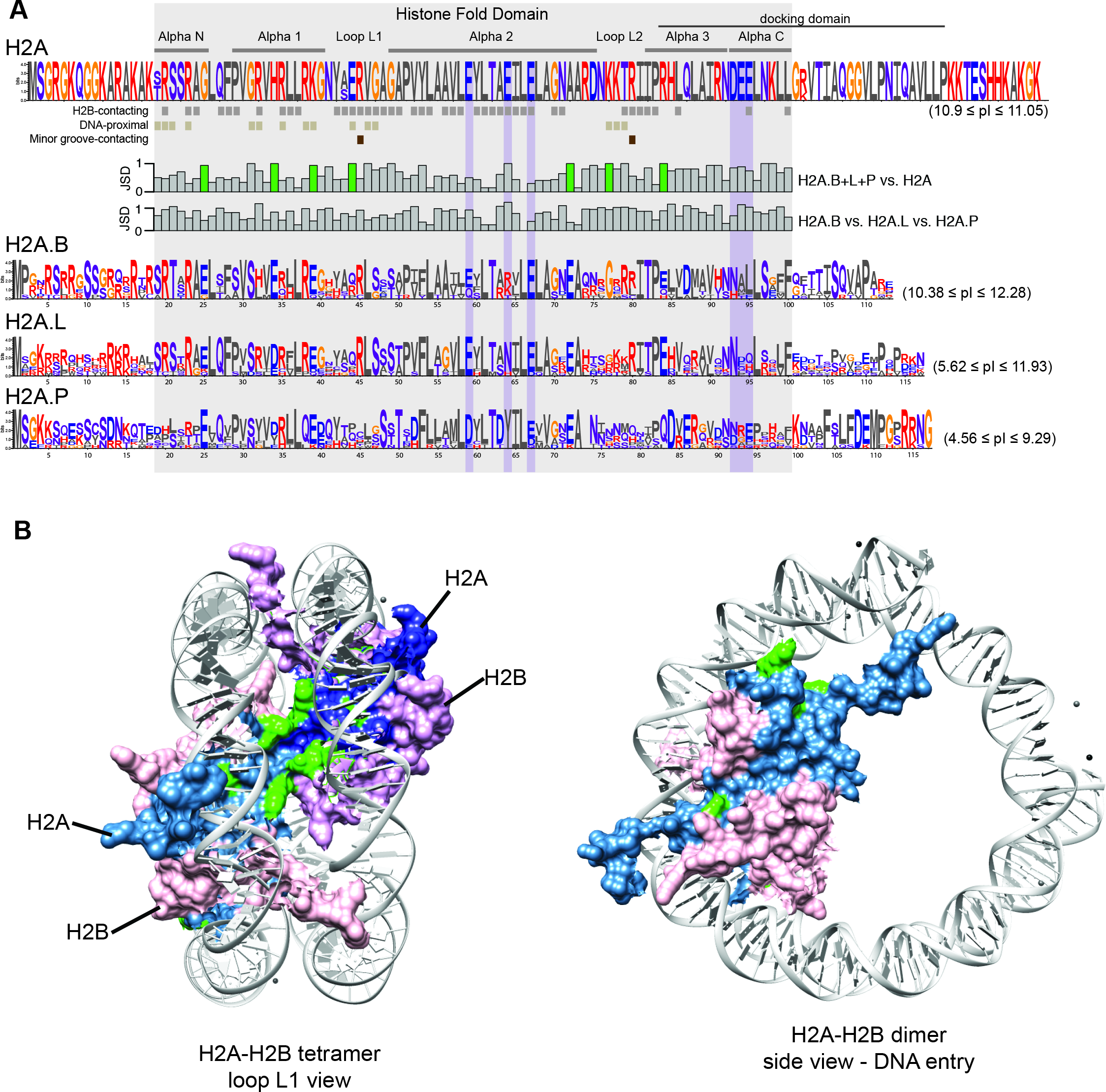
Short H2A variants have distinct protein features. **(A)** Logo plots of protein alignments of canonical and three of four short H2As (H2A.B, H2A.P and H2A.L) across an identical set of representative placental mammals (see methods). Residues are colored based on their biochemical proprieties; hydrophobic residues in grey; positively charged in red; negatively charged in blue; polar in purple and others (Gly and Cys) in orange. Residues in proximity to H2B (grey squares, at least 4Å), DNA (light brown, 5Å) or buried in the minor groove (dark brown) are annotated below the H2A logo. The histone fold domain and acidic patch are boxed in grey and purple respectively. Jensen-Shannon distances (JSD) measured at each amino acid position between combinations of logos are indicated as bar graphs (ranging from 0, very similar, to 1, very different). Charge-altering changes that are conserved among short H2As but different from canonical H2A are highlighted using green-colored bars. The average isoelectric point (pI) across the eight representative mammals are shown in parentheses. **(B)** Left panel, loop L1 view of a H2A-H2B tetramer in the context of the nucleosome crystal structure (Luger et al., 1997, see methods). Right panel, H2A-H2B dimer view from the DNA entry site. H2A molecule surfaces are shown in light and dark blue, H2B in pink and purple. DNA is shown in grey. Charge-altering discriminative residues highlighted in panel (A) are mapped in green onto the H2A structure.

Consistent with previous findings, our analysis indicates that many key structural residues of the HFD are conserved between canonical H2As and short histone H2A variants. However, our analysis also identifies features that differ between all short H2A variants and canonical H2A (Figure 4A). In order to highlight amino acid positions in the histone fold domain that distinguish short H2A variants from canonical H2A, we used the Jensen-Shannon distance (JSD) to compare the aligned sequences (Figure 4A bar graphs). We measured the JSD between the three analyzed short H2As and canonical H2A, to find positions that are shared by all short variants but are universally different from canonical H2A (upper graph). Represented by JSD values close to 1, many residues within the alpha-helices and the loops of the HFD differ between short histone variants and canonical H2A but are conserved across the three short H2As examined. The most obvious of these differences have been noted previously (Gautier, et al. 2004; Eirin-Lopez, et al. 2008; Gonzalez-Romero, et al. 2008; Ishibashi, et al. 2010; Bonisch and Hake 2012; Talbert, et al. 2012; Rivera-Casas, et al. 2016; Buschbeck and Hake 2017). First, the C-terminal tail of all three variants is shorter than in canonical H2A. Second, short histone H2A variants no longer conserve the acidic patch that is involved in nucleosome-nucleosome interactions (Luger, et al. 1997; Chakravarthy, et al. 2004). The high divergence of Loop1 (or loop L1) in short histone H2A variants compared to canonical H2A proteins could affect overall stability of the H2A::H2A interaction within each octameric nucleosome and therefore the nucleosome itself (Luger, et al. 1997; Chakravarthy, et al. 2004). We hypothesize that these alterations in the HFD, especially changes in loop L1, make it increasingly unlikely that short histone H2A variants would be able to form stable heterotypic dimers with canonical H2A. In addition, we identified 7 discriminative positions within the HFD that are charge-altering changes common to all short H2As. In almost all cases, short H2As preserve acidic residues at positions spread over the entire HFD in contrast to uncharged or basic residues in canonical H2A (Figure 4A). Mapping these charge altering residues (mostly acidic) onto the published nucleosome crystal structure (Luger, et al. 1997) shows that they cluster around H2A::H2A contact sites in proximity to DNA (Figure 4B). Together with previous reports of the shorter wrap of DNA around nucleosomes containing short H2A variants (Bao, et al. 2004; Doyen, et al. 2006; Syed, et al. 2009), our findings provide further evidence that sequence features of the short histone H2A variants may predispose them to form nucleosomes with lower stability and packing density.

Using a similar methodology, we also calculated JSD between all three analyzed short histone H2A variants, to find positions that are conserved within each of the three variants but that distinguish them from one another (lower graph), suggesting further differentiation after their common ancestry (Figure 4A, JSD H2A.B vs H2A.L vs H2A.P). For example, we note that H2A.P has acquired several additional acidic changes after divergence from the other two short histone H2A variants (E8, D13, D/E40, D52, D63, D84, E86, D108 and E109), including sites predicted to contact DNA and H2B. Furthermore, H2A.P has lost two arginine (R) residues in loop 1 and 2 that contact the DNA minor-groove. These arginine residues are conserved in canonical H2A, as well as including H2A.B and H2A.L, and their loss might further reduce the stability of H2A.P-containing nucleosomes. In fact, changes in H2A.P have been so extensive that the theoretical isoelectric point (pI) of full-length H2A.P averages 5.5 across the eight species used, which is dramatically different from the pI of canonical H2A, at 11.0, or that of the other short histone H2A variants (11.51 for H2A.B and 9.83 for H2A.L, Figure 4A). These changes in acidity would result in especially destabilized histone/DNA interactions for H2A.P. Indeed, such an acidic pI is highly unexpected for a DNA-packaging protein.

While most previous analyses focused on the HFD of short histone variants, we also found key features of short H2A histone tails that further differentiate them from canonical H2A. The N-and C-terminal tails of H2A.B, H2A.L, and H2A.P cannot be well aligned to either canonical H2A or to each another. Despite this divergence, the tails still show some functional constraint within each variant type (Figure 4A), indicating that these regions may carry out important roles that differ between variants. In fact, more conserved features distinguishing H2A.B, H2A.L and H2A.P from one another appear in the N-and C-terminal tails than in the HFD. This is particularly evident for H2A.P, where the last 14 amino acids appear to be more constrained than the rest of the protein. These distinguishing features would suggest that each of the short histone H2A variants interact with different non-histone proteins via their tails.

### H2A.Q: a novel short histone variant

Our finding of H2A.Q highlights both the difficulty in fully annotating rapidly evolving histone variants as well as the utility of a phylogenomic framework to identify and annotate new members. In this instance, our framework first identified *H2A.Q* as a gene that was present in a shared syntenic location in many mammalian genomes, but did not overlap with any of the well annotated H2A.L loci. Moreover, phylogenetic analyses clearly implicated it as a divergent clade of histones, more similar to *H2A.P* than to *H2A.L*. H2A.Q has an N-terminal tail of typical size, but a shorter C-terminal tail than other short histone H2A variants (Figure 5A). H2A.Q shares many of the features that discriminate short histone H2A variants H2A.B, H2A.L, and H2A.P from canonical H2A, including the loss of the acidic patch and histone-docking domain (Figure 4). Just like the other short histone variants, H2A.Q is also missing the two DNA minor groove-contacting arginine residues in Loop1 and Loop2 present in canonical H2A (bold red, Figure 5A). Similarly, H2A.Q is different from canonical H2A in the seven amino acid positions (bold black, Figure 5A) that distinguish short histone H2A variants from canonical H2A.

**Figure 5:**
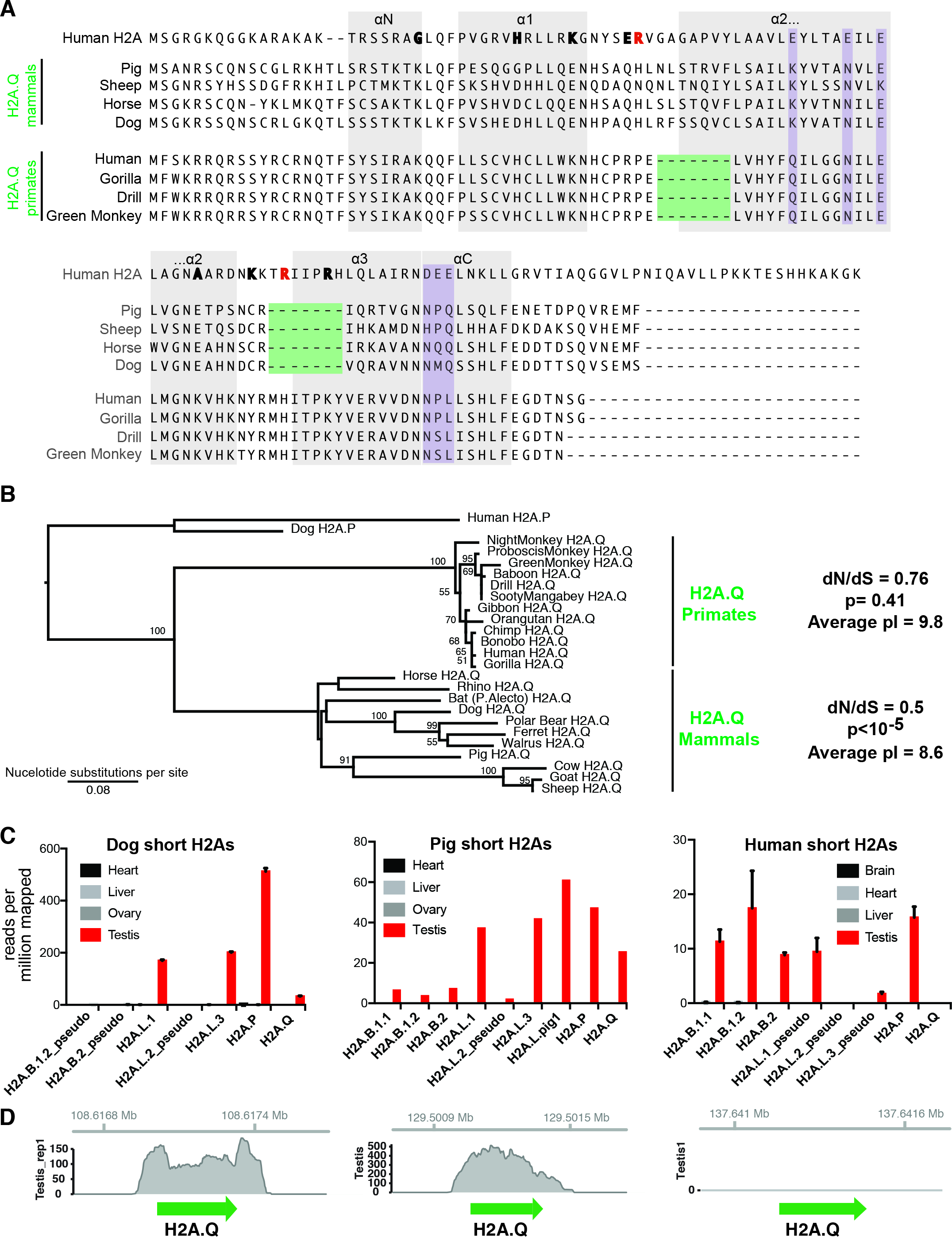
The novel short histone H2A variant H2A.Q. **(A)** An alignment of representative primate and non-primate H2A.Q proteins, along with canonical H2A. Charge-altering residues that distinguish short histone variants from canonical H2A (see Figure 4) are highlighted in bold black and the two arginine residues that make minor groove contacts are highlighted in bold red; the residues comprising the acidic patch are shown using purple shading. **(B)** Maximum likelihood nucleotide phylogeny of mammalian *H2A.Q* genes. Human and pig *H2A.P* are used as outgroup sequences. Note the much shorter branch lengths of the simian primate *H2A.Q* genes compared to those from non-primate mammals, despite the fact that the latter have dN/dS values more consistent with a protein-coding open reading frame. **(C)** RNA-sequencing analyses of short H2As from selected somatic and germline tissues of dog, pig and human shows that *H2A.Q*, like other short H2As, shows testis-specific expression in dog and pig. *H2A.Q* does not appear to be transcribed in human testis, supporting evolutionary analysis that suggests it may now be non-functional in primates. Error bars show standard deviation of biological replicates, where available. **(D)** Coverage plots provide additional support for robust transcription of *H2A.Q* in dog and pig testis.

Atypically, however, H2A.Q appears to have undergone distinct deletions in its HFD. Simian primates have a 7 amino acid deletion removing parts of Loop 1 and the alpha2 helix, whereas non-primate mammals have a distinct 7 aa deletion that removes parts of Loop 2 and alpha helix3 (Figure 5A). While no extant intact H2A.Q gene exists that lacks either deletion, *H2A.Q* pseudogenes from lemur genomes support the existence of an ancestral full-length *H2A.Q* gene that lacked both deletions, confirming that the two deletions were independently acquired (Supp. Figure 3). Either of these deletions would dramatically affect the topology of both the H2A.Q HFD as well as any nucleosome structure that uses H2A.Q instead of any other canonical or variant H2A protein.

While H2A.Q is clearly a pseudogene in several clades including rodents and felines, with premature stop codons and/or frameshifts, it appears to encode a complete ORF in many other species. This observation, and the atypical protein features, prompted us to investigate whether H2A.Q orthologs are functional genes or if they might be drifting towards becoming non-functional pseudogenes. We constructed a nucleotide phylogeny of a broad set of potentially coding H2A.Qs (Figure 5B). This phylogeny confirmed the orthology of *H2A.Q* genes and showed that non-primate H2A.Q sequences are diversifying more rapidly than simian primate sequences (i.e. comparing branch length between dog and polar bear, ~47My of divergence, and between night monkey and human, ~42My). In addition, this allowed us to estimate dN/dS to assess whether purifying selection acts on H2A.Q. We find that the overall dN/dS for non-primate *H2A.Q* genes is 0.5, supporting its conservation as a protein-coding gene (p<10^-5^ rejecting a model of neutral evolution). However, we cannot reject neutral evolution for the primate *H2A.Q* orthologs (dN/dS =0.76, p =0.41); this may suggest that H2A.Q is non-functional in primates. Furthermore, closer examination of the simian primate H2A.Q genes to their pig and lemur (pseudogene) orthologs (Supp. Figure 3) revealed that a short region of human and other primate H2A.Q genes may be frameshifted relative to pig and other intact H2A.Qs: two nearby compensating frameshift changes make it difficult to determine whether the reading frame remains intact or not (Supp. Figure 3).

In order to further explore the functional potential of H2A.Q, we analyzed publically available RNA-seq datasets to determine whether *H2A.Q* is transcribed (Figure 5C, Supp. Table 3). We find excellent support for transcription of dog and pig *H2A.Q* genes, which appear to be testis-specific like the other short H2A variants. However, we are unable to detect *H2A.Q* expression in any tissue in humans (Figure 5C) or several other primates (not shown). Taken together, our analyses supports the idea that *H2A.Q* is functional and subject to accelerated evolution in some species, but that it is no longer functional in primates.

### Features of “proto-short H2As”: mmH2As in marsupials and monotremes

Our histone H2A surveys also revealed five previously undescribed H2A variants in monotreme and marsupial genomes, referred to as mmH2As. Based on our phylogenetic analysis, we inferred that mmH2A variants represent the most closely related histones to the short H2A variants of eutherian mammals. These findings suggest that a “proto-short H2A variant” originated early during mammalian evolution. Several protein features of mmH2A variants appear to be intermediate between short histone H2A variants and canonical H2A. Similar to canonical H2A, mmH2A variants appear conserved, as evident from alignments and logo plots (Figure 6A, 6B) and from short branch lengths in the phylogeny (Figure 3). They are also more closely related to canonical H2A at most residues within the HFD and at the docking domain (Figure 6A). Interestingly, in contrast to the shorter C-termini of short histone H2A variants, mmH2A have extended C-termini compared to canonical H2As.

**Figure 6:**
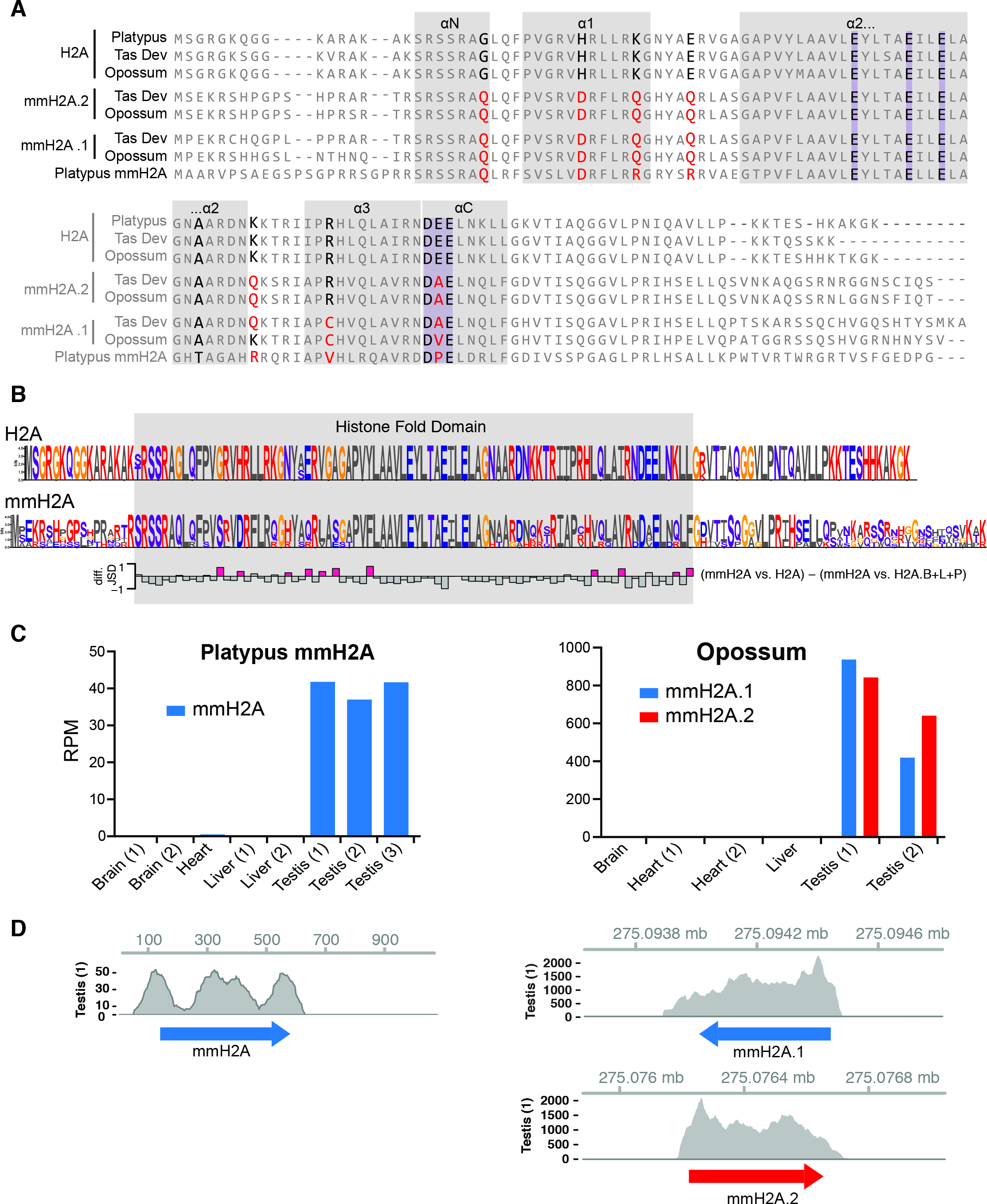
mmH2A variants in basally-branching mammalian lineages. **(A)** A protein alignment of all five sequenced mmH2A variants along with canonical H2A representatives. We highlight the acidic patch residues in purple boxes (note the deviation from the DEE consensus in alphaC). We also highlight in boldface the seven positions that distinguish short histone variants from canonical H2A (Figure 4); charge-altering residues in mmH2A variants at these positions are highlighted in bold red. **(B)** Protein logos of mmH2A variants and canonical H2A further highlight that they share several features in common to the exclusion of short histone H2A variants. Jensen-Shannon distance (JSD) differences show that overall, mmH2As are more like canonical H2A than short H2As, but that there are a few residues, highlighted in pink, where mmH2As are more similar to short histone H2As than canonical H2A. **(C)** RNA-seq analyses of platypus and opossum somatic and germline tissues shows that the mmH2A genes have testis-specific expression. Numbers in parenthesis represent individual biological replicates. **(D)** Coverage plots provide additional support for robust transcription of *mmH2A* in platypus and opossum.

To reveal mmH2A features that were shared with short H2As, we computed the difference in JSD between mmH2A vs. canonical H2A and mmH2A vs. all short H2As (Figure 6B bottom bar graph). While overall, mmH2A appears closer to canonical H2A than to short H2A variants, as evidenced by negative JSD differences, there are several positions at which mmH2A more closely resembles short H2As than canonical H2A (pink bars). These include residues at the N-term and C-term residues in of the HFD, a few of which exhibit charge-altering changes that might alter histone/DNA interactions. we also observe a partial loss of the acidic patch (DEE motif in the alphaC helix, Figure 6A). These features likely reflect the “intermediate” evolutionary position of mmH2A with respect to canonical H2A and the short H2A variants.

Finally, we analyzed publically available RNA-seq datasets from platypus and opossum and found that, like short H2A variants, mmH2A variants appear to be testis-specific (Figure 6C). Thus, marsupial and monotreme variants likely fixed a “transitional stage” in the mammalian-specific evolution of testis short H2A variants. Their functional characterization may reveal important insights into the evolution of this distinct class of histone variants. Nevertheless, the discovery that these variants are not “short” H2As, by definition, conclusively demonstrates the origin of short H2A variants *per se* in the common ancestor of all eutherian mammals.

### Concerted evolution of H2A.B and H2A.L gene duplicates

Although we initially expected to see that each ancestral H2A variant locus would diverge gradually from the other loci over time, our phylogenetic analysis revealed many instances where individual *H2A.B* or *H2A.L* genes within species shared more sequence similarity to each other than to their “true orthologs”, as defined by synteny (Figure 2, Figure 3). This pattern of evolution we see for *H2A.B* and *H2A.L* indicates that these genes could be subject to concerted evolution in some species, whereby there is selective pressure to keep multiple gene copies within a species highly homogeneous. Likewise, canonical histone loci dispersed across mammalian genomes are highly homogeneous within species (Coen, et al. 1982; Rooney, et al. 2000; Marzluff, et al. 2002; Nei and Rooney 2005; Marzluff, et al. 2008). For canonical histones, the evolutionary mechanism maintaining homogeneity appears to be a process of gene duplication and loss (birth-and-death) combined with strong purifying selection at the protein sequence level (Rooney, et al. 2002). In contrast, we observe sequence homogenization between short H2As that have resided in the same syntenic region for over 100 My, which strongly suggests that gene conversion is “overwriting” pre-existing loci.

To further test the hypothesis of gene conversion, we analyzed nucleotide sequences to explore the evolutionary path that led to sequence homogeneity within species. We first examined primate *H2A.B* genes (Figure 7A). We collected and aligned all *H2A.B-*related DNA sequences from the genomes of six primates representing roughly 40 million years of evolution. In addition to the homologs at the three ancestral H2A.B loci (*H2A.B.1.1*, *H2A.B.1.2* and *H2A.B.2*), we discovered additional genes in marmoset in non-syntenic loci (*H2A.B.Mar1 to 4*). We performed GARD analysis (Kosakovsky Pond, et al. 2006) to search for internal recombination breakpoints, but found none. We then estimated a maximum-likelihood phylogeny using pig *H2A.B* genes as an outgroup (Figure 7A). This phylogeny clearly groups *H2A.B* duplicates by species rather than by syntenic genomic location. In most cases, *H2A.B* duplicates are 100% identical within species at the nucleotide level.

**Figure 7:**
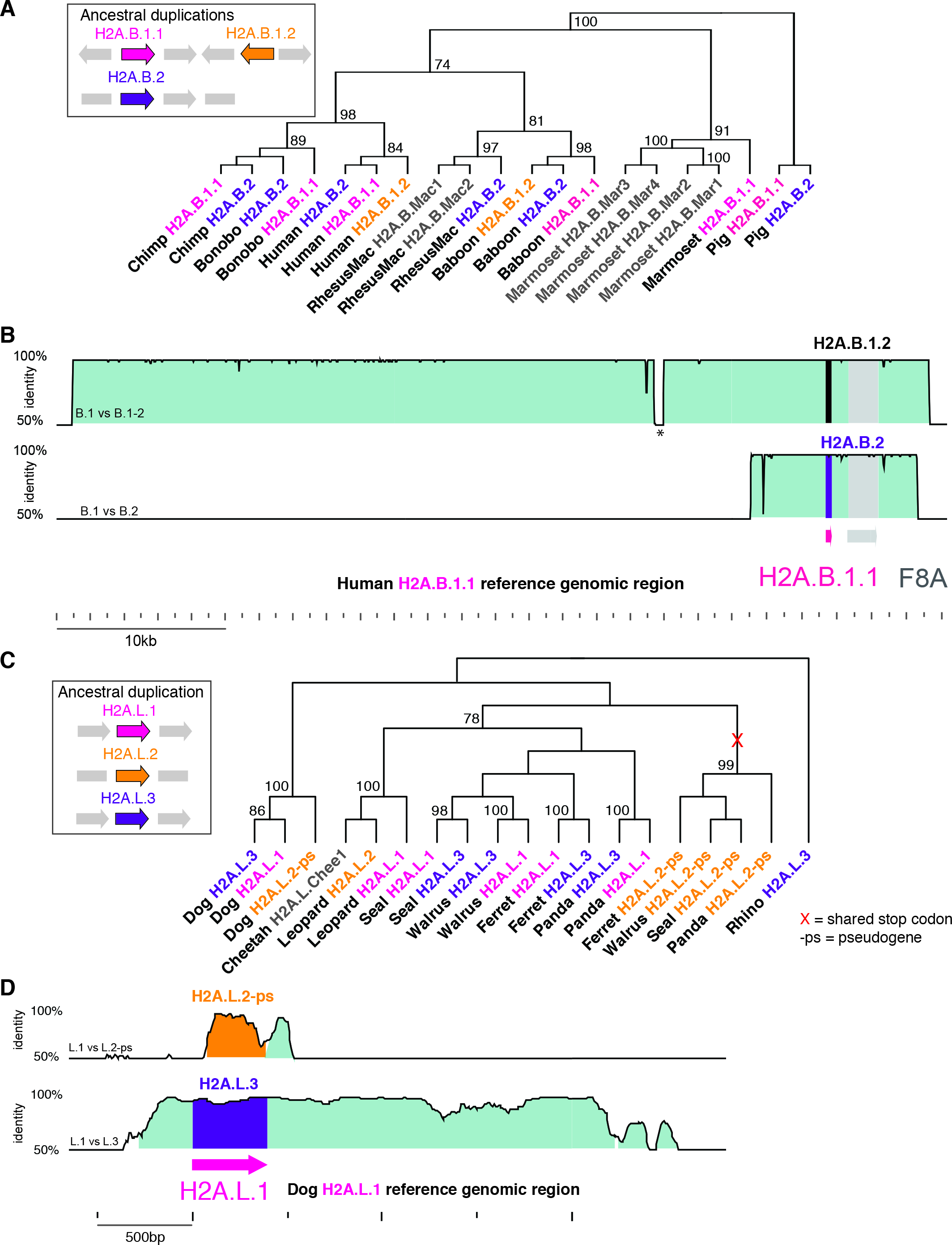
Concerted evolution by gene conversion among primate *H2A.B* and carnivore *H2A.L* genes. **(A)** Maximum-likelihood nucleotide phylogeny of representative primate H2A.B genes rooted using pig outgroups. Bootstrap values are shown for nodes with >50% support. A cartoon shows the loci containing ancestral duplications *H2A.B.1.1* (pink), *H2A.B.1.2* (orange) and *H2A.B.2* (purple), and gene names are colored accordingly in the tree. We also include several marmoset-specific duplicate genes at non-syntenic loci (denoted “_Mar1 to 4”). **(B)** VISTA plots showing very high nucleotide identity across large genomic regions flanking the three human *H2A.B* genes, comparing *H2A.B.1.1* genomic sequence as a reference to *H2A.B.1.2* (upper plot) and *H2A.B.2* (lower plot). An interruption in high identity (asterisk) is due to a recent transposon insertion. **(C)** Like panel A, but for carnivore *H2A.L* genes, using rhinoceros as an outgroup. The *H2A.L.2* locus acquired a premature stop codon in the common ancestor of ferret, walrus, seal and panda, so is a pseudogene ("-ps") in those extant species, and appears to have been exempt from gene conversion since its loss of function. Dog *H2A.L.2* is also a pseudogene but its function was likely lost independently. **(D)** VISTA plots for dog *H2A.L* genes using *H2A.L.1* as reference. Gene conversion appears to have homogenized a ~2.5kb stretch between *H2A.L.1* and *H2A.L.3* but does not appear to have affected the surrounding sequences of the *H2A.L.2* pseudogene.

Using VISTA plots (Figure 7B) to expand our analysis and include neighboring sequences, we found longer regions of high homology between species. Between human *H2A.B.1.1* and *H2A.B.1.2*, a nearly 50 kb region is almost identical with sharp drop-offs in identity on either side. Human *H2A.B.1.1* and *H2A.B.2* share a shorter nearly identical region (about 10 kb). These results suggest that *H2A.B* duplicates were indeed subject to very recent gene conversion, which has occurred recurrently in the genomes of multiple species. Large gene conversion tracts that include not only the coding sequences but also the neighboring sequence likely drove this sequence homogenization. This is a significant contrast to the birth-and-death mechanism of concerted evolution in canonical histone genes, which show homogeneous protein sequences but display many synonymous nucleotide changes (Rooney, et al. 2002). The gene conversion we observe among the three genes is surprising given the relatively large distance between the B1 and B2 loci (576 kb) and since the frequency of gene conversion is inversely proportional to the genomic distance between loci (Schildkraut, et al. 2005).

We next investigated whether gene conversion could also be seen in *H2A.L* genes. Since *H2A.L* genes are likely to be pseudogenes in many primates (see above), we instead investigated sequenced carnivore genomes to look for signs of gene conversion in *H2A.L* genes. Similar to primate *H2A.B* genes, we observed similar signals of gene conversion in carnivore *H2A.L*s, including between paralogs that are surprisingly far apart in the genome (1.06 Mb between dog *H2A.L.1* and *H2A.L.3*) (Figure 7C and 7D). However, not all carnivore *H2A.L* loci participate in this concerted evolution. Following the acquisition of a premature stop codon in the common ancestor of ferret, walrus, seal and panda, the *H2A.L.2* locus appears to have been exempt from sequence homogenization. We find several other examples where a locus is no longer involved in gene conversion after the intact ORF is lost. For example, the *H2A.L.2* locus acquired a stop codon in the ancestor of all primates and has retained an independent evolutionary path *i.e.*, no longer subject to gene conversion since that time (Supp. Figure 1). Furthermore, the *H2A.L.1* and *H2A.L.3* loci described above as pseudogenes in catarrhine primates also evolve independently, while the still-functional New World monkey *H2A.L* genes are mostly homogeneous within species (Supp. Figure 1). Thus, like *H2A.B*, we see a recurrent but sporadic pattern of gene conversion for *H2A.L* genes.

The mouse *H2A.L* genes already seem exceptional among short H2A variants in their greatly expanded copy number (Ferguson, et al. 2009). We wondered whether these new *H2A.L* paralogs also evolve by concerted evolution or whether some copies might have escaped homogenization. We collected *H2A.L*-like sequences from a number of additional rodent genomes, and created a phylogeny (Supp. Figure 2). We found that many of the reported mouse *H2A.L* duplicates, including the chromosome 2 copy shown to be important for protamine deposition (Barral, et al. 2017), predate the divergence of the *Mus* species. However, our analysis did not find any evidence of ongoing concerted evolution; each of the ancestrally duplicated loci, *H2A.L.1, H2A.L.2* and *H2A.L.3*, groups separately on the phylogeny, and some of the recent duplicates (including copies on chromosome 2 and Y chromosome copies) also appear to have their own evolutionary trajectories. While this loss of gene conversion can be explained by loss of function for the *H2A.L.1* locus, which contains a pseudogene in most rodent genomes, most genes in the *H2A.L.3* and *H2A.L.2* loci still appear functional, as do the recently duplicated copies. We suggest that the lack of concerted evolution reflects ongoing diversification and functional specialization of subgroups of rodent *H2A.L* genes. This hypothesis is supported by a recent finding that knocking out just a single mouse *H2A.L.chr2* gene renders male mice infertile (Barral, et al. 2017); other *H2A.L* duplicates cannot compensate for loss of *H2A.L.chr2*.

Overall, we conclude that concerted evolution has played a significant role in shaping the evolution of active *H2A.B* and *H2A.L* genes in some but not all eutherian mammal genomes. This mode of evolution appears distinct from the previously described birth-and-death model that is applicable to histone multigene families in animal genomes (Piontkivska, et al. 2002; Rooney, et al. 2002).

### Accelerated evolution and diversifying selection of short H2A variants

Both the phylogenetic analysis (Figure 3) and our protein alignments (Figure 4) suggest that short H2A variants are subject to much lower evolutionary constraints than canonical histones or indeed, other histone H2A variants. To directly test this hypothesis, we compared divergence levels between different histone H2A variants (the four short H2A variants, mmH2A, canonical H2A, H2A.X, H2A.Z, macroH2A and testis-specific H2A.1), the cenH3 variant that is rapidly evolving (Malik and Henikoff 2001; Talbert, et al. 2002), and a testis-specific H2B variant, H2B.1. We adopted two analyses to evaluate whether the high divergence of short histone H2A variants is unusual.

First, we estimated the dN/dS along multiple codon alignments of each variant across exactly the same eight species used for our previous logo plot analyses (Figure 4, Methods). We used PAML’s codeml with NSsites model 0 that assumes a single evolutionary rate across all sites and species. We obtained overall dN/dS values of 0.24 for *H2A.B*, 0.32 for *H2A.L* and 0.58 for *H2A.P*, compared to 0.003 for canonical H2A. In each case, PAML provides very strong evidence that the overall selective regime acting on these ORFs is one of purifying selection, rather than neutral evolution (M0 vs M0 with dN/dS fixed at 1), supporting the idea that they are functional variants that contribute to mammalian fitness on some level. This is especially interesting in the case of H2A.P for which RNA expression but no protein expression has been observed. Our dN/dS analysis allows us to conclusively reject the model of neutral evolution (p<10^-4^) suggesting that H2A.P indeed must encode a protein. As previously discussed (Figure 5B), when we similarly analyzed dN/dS values for non-primate or simian primate *H2A.Q*, we obtained strong evidence of purifying selection for the former (dN/dS = 0.5), but not the latter (dN/dS =0.76). Thus, although we can conclusively reject neutral evolution for H2A.Q in non-primate mammals (see Figure 5B), we cannot for simian primates, in spite of the fact that we detected no obvious disruptions of open reading frame.

In an orthogonal method, we performed whole protein pairwise comparisons of a human copy of each histone variant to a series of orthologs from species of increasing evolutionary divergence. This allows us to examine the rate of protein evolution at different ages of divergence and over long evolutionary times. Unfortunately, since *H2A.L* genes are pseudogenized in human, we instead used dog *H2A.L* as the reference sequence for this variant. Similarly, given the uncertainty regarding *H2A.Q* genes in primates, we focused on non-primate comparisons for *H2A.Q* as well. We also examined divergence between mmH2A variants that are the closest relatives of the short H2A variants. We plotted amino acid identity as a function of species divergence time according to TimeTree (Figure 8A) (Hedges, et al. 2015). As previously mentioned, our analysis reveals very high conservation of canonical H2A and most other variant histones, including testis-specific H2A.1 and H2B.1, and mmH2A variants. By contrast, cenH3 shows greater cross-species divergence than canonical H2A, as expected given its rapid evolution. We found that the short H2A variants are also diverging more rapidly than their canonical counterpart suggesting that they may be subject to accelerated evolution like cenH3.

**Figure 8:**
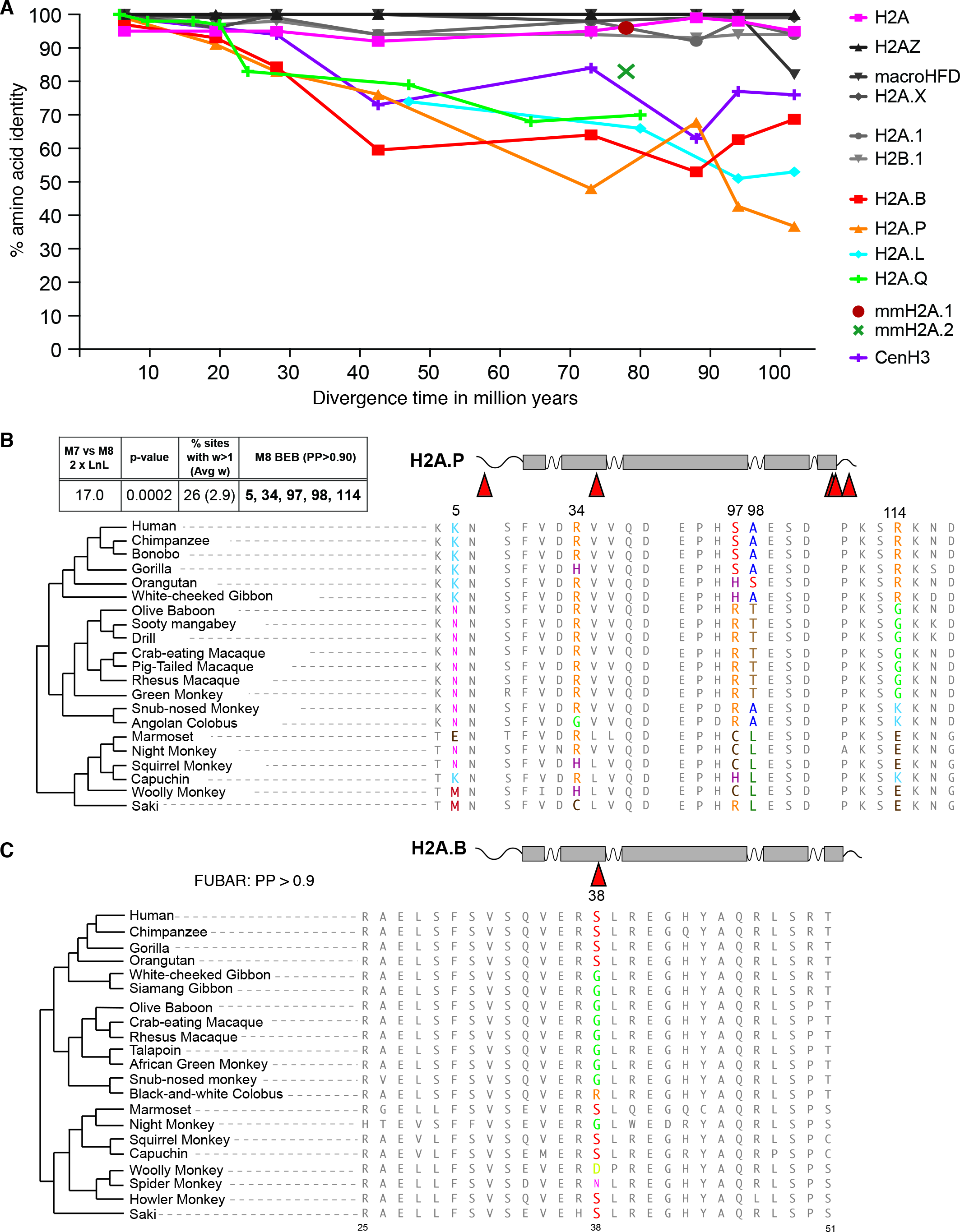
Rapid evolution of short H2As. **(A)** Pairwise amino-acid identities (y-axis) between pairs of mammalian H2A genes as a function of species divergence time (x-axis, see methods). Marsupial variants 1 and 2 are shown as single points, since each is represented by only two species diverged by 78 million years. **(B)** Portions of alignments surrounding positively selected sites in simian primate H2A.P proteins (PAML, top left box). **(C)** A portion of the alignment of simian primate H2A.B proteins showing a positively selected residue at site 38 (colored amino acids), identified by FUBAR.

Since none of the four short H2A variants are universally conserved, it is possible that the accelerated divergence we observe in *H2A.B*, *H2A.L*, *H2A.P* and *H2A.Q* is the result of relaxed purifying selection. Alternatively, the high divergence could reflect diversifying selection, possibly with even greater strength than was experienced by cenH3 or any other histone variant. To search for diversifying selection, we collected presumed intact *H2A.B, H2A.P* and *H2A.Q* ortholog sequences from simian primates, which exhibit an ideal evolutionary divergence for analyses of selective pressures. In each case, we collected sequences from databases as well as by PCR amplification and Sanger sequencing. For *H2A.B*, we used a single duplicate from each species, owing to the signature of gene conversion. We codon-aligned sequences and used maximum-likelihood approaches implemented in PAML (Yang 1997) and HyPhy (Pond, et al. 2005) packages to compare the rates of non-synonymous (dN) to synonymous (dS) substitutions. We obtained identical results from both a phylogeny inferred from each alignment and the published species tree (Perelman, et al. 2011).

We detected clear evidence of diversifying selection in *H2A.P* in simian primates. Our analysis used a total of 21 *H2A.P* sequences, including 19 from assembled genomes, and 2 obtained by PCR and sequencing (Methods, Supp. Data 3). PAML analysis (Figure 6B) reveals diversifying selection (M7 vs. M8, p-value < 0.001) on 26% of the sites, with average dN/dS of 2.9. Five sites (codon 5, 34, 97, 98 and 114) had high posterior probabilities of evolving under diversifying selection (>0.9, M8 BEB) with several others just under that threshold. Codon 5 is in the N-terminal tail that is known to play a role in the post-translational modification of histone proteins. Codon 34 is in helix 1 and is predicted to be proximal to both DNA and H2B. Codons 97 and 98 are in the extension of alpha-helix 3 (“alpha-C”) and codon 114 is located in the unstructured C-term tail. FUBAR analysis (HyPhy package (Murrell, et al. 2013)) confirms positive selection on site 97 with additional evidence at codons 17, 43, 79 and 116 (posterior probability >0.9). Additional methods to detect positive selection in the HyPhy package (REL, MEME) add further support to our finding of positive selection (Murrell, et al. 2012; Smith, et al. 2015). The evidence for diversifying selection of *H2A.P* is comparable to that previously obtained for *cenH3* (*CENP-A*) orthologs in primates (Schueler, et al. 2010).

In contrast to *H2A.P*, we found no evidence of positive selection acting on *H2A.Q* in either simian primates or Cetartiodactyla, despite high overall dN/dS values (Suppl. Table 2). The evidence for diversifying selection is more equivocal for *H2A.B*. Our analysis used a total of 21 *H2A.B* sequences, including 14 from assembled genomes and 7 obtained by PCR and sequencing (Methods, Supp. Data 3). FUBAR finds a single site, codon 38, with strong evidence for diversifying selection (posterior probability >0.9, Figure 6C). This site, found in alpha-helix 1 of the HFD, is also among those highlighted by the FEL, REL and MEME algorithms as being under positive selection. PAML finds evidence for positive selection using one set of starting parameters; however, statistical tests do not reach conventional thresholds for significance and the finding of diversifying selection is not robust to the use of alternative codon models.

Overall, our results indicate that *H2A.P* and possibly *H2A.B* have been subject to diversifying selection in simian primates, which could partly explain the greater divergence of short H2A histone variants compared to other H2A histones in mammals (Figure 8A).

## Discussion

In this study, we describe the origin and evolutionary dynamics of four rapidly evolving H2A variants. We infer that a precursor to these short H2A variants arose in the common ancestor of all eutherian mammals. Subsequently, the C-terminal region was truncated, and the variant duplicated and diversified into the four “types” of short H2A variants we see today: H2A.B, H2A.L, H2A.P and H2A.Q. Based on shared synteny, we infer that the last common eutherian mammal ancestor encoded eight short H2A variant genes on the X chromosome: 3 *H2A.B* genes, 3 *H2A.L* genes, a single *H2A.P* gene and a single *H2A.Q* gene. However, the repertoire of short H2A histone variants has since been remarkably plastic in different lineages, with some species entirely losing functional *H2A.B, H2A.P, H2A.Q* or *H2A.L* genes, and other lineages experiencing additional duplications. *Mus musculus* provides a particularly dramatic example of lineage-specific duplication, with 18 functional *H2A.L* loci and 2 pseudogenes (Ferguson, et al. 2009). The finding that no single short H2A variant is universally indispensable to all mammals might suggest that they are functionally non-essential or redundant with each other. However, the finding that deletion of even a single *H2A.L.chr2* gene in mice leads to male infertility renders both these possibilities unlikely (Barral, et al. 2017). Alternatively, the short histone variants might have variable functions between different mammals, whereby different short histone variants may have taken on similar functions in different lineages. Indeed, H2A.B and H2A.L may perform similar functions in mice and humans, despite their evolutionary divergence. Mouse H2A.B is primarily expressed during and after meiosis but is not present on sperm chromatin, which contains H2A.L instead. However, in humans, *H2A.L* appears to have pseudogenized and sperm chromatin now contains H2A.B (Baker, et al. 2007; Baker, et al. 2008; El Kennani, et al. 2017).

In addition to identifying a novel fourth short histone variant, H2A.Q, our analysis also clarifies the relationships amongst the four short H2A variants, which have previously been confused with each other due to lack of comprehensive phylogenetic analyses and inconsistent nomenclature. Indeed, we firmly establish that *H2A.B*, *H2A.L*, *H2A.P* and *H2AQ* represent four related but distinct evolutionary clades. Furthermore, although H2A.P protein has never been detected in mouse or human germ cells (Baker, et al. 2007; Baker, et al. 2008; El Kennani, et al. 2017), our analysis strongly argues that *H2A.P* is indeed a functional protein-coding gene, as is *H2A.Q*, at least in non-primate mammals. Failure to detect H2A.P protein in testis samples may be due to expression in a limited subset of cells, or perhaps even because *H2A.P* mRNAs, which accumulate during spermatogenesis, may actually be delivered to the embryo and translated following fertilization.

Our work also provides a more comprehensive understanding of the amino acid constraints that likely drove the diversification and specialization of short H2A variants. Consistent with previous analyses, we find that four short H2A variants have amino acid changes that suggest they are “short-wrappers”, and lack the acidic patch responsible for tight chromatin packaging (Luger, et al. 1997; Chakravarthy, et al. 2004). Additionally, we find that all short histone variants differ from canonical H2A at some key positions, many of which are in or near loop L1 of the HFD. Such changes may therefore affect H2A::H2A interactions within the nucleosome and may preclude formation of heterotypic nucleosomes that contain both canonical and short variant H2A proteins.

Although H2A.B, H2A.P, H2A.Q and H2A.L might have independent functions, as suggested by their independent evolutionary trajectories, we find very few conserved residues in the HFD that differentiate each variant type. However, we find evidence of specialization in the N-and C-terminal tails of these variants, especially in H2A.P and H2A.B. We therefore predict that the short H2A variants may dramatically differ in their protein-protein interactions. These features suggest that rapid specialization occurred after the separation of short histone H2A variants into four types. And yet, none of the short histone variants are universally conserved in eutherian mammals (Figure 2). It is unclear whether this is due to altered requirements for short histone variants across male germlines of different eutherian mammals, or due to some degree of functional redundancy. Elucidation of the molecular functions of short H2As will help distinguish between these possibilities.

Our analysis also revealed strong evidence of concerted evolution among the multiple *H2A.B* and *H2A.L* paralogs in multiple mammalian genomes. We find that recurrent gene conversion overwrites sequences that have resided in ancestrally syntenic loci. Although canonical histone gene duplicates are also highly homogeneous within species, this likely occurs by a distinct evolutionary mechanism: a combination of birth-and-death evolution and purifying selection at the protein sequence level (Rooney, et al. 2002; Nei and Rooney 2005). However, rodent *H2A.L* duplicates follow independent evolutionary trajectories, with dramatic expansion of *H2A.L.2*-like paralogs in mouse. The lack of gene conversion among rodent *H2A.L* genes is consistent with the hypothesis that they have acquired non-redundant functions, supported by the deletion of *H2A.L.chr2* (Barral, et al. 2017). Our findings also suggest that, at least in some instances, the different paralogs of short histone variants within a species (e.g., H2A.B in mouse) are unlikely to encode dramatically different functions (Soboleva, et al. 2017). Furthermore, there would be little point in ascribing specific functions to orthologs of individual short histone variants, as orthology would not be indicative of sequence evolution or of function.

We find that short H2A variants show greater evolutionary divergence between species than even cenH3, the fastest-evolving histone variant examined to date (Malik and Henikoff 2001; Talbert, et al. 2002). Rapid divergence of short histone variants could be partly the result of decreased functional constraint due to their male germline specificity. However, in at least one lineage, primates, this divergence is partly due to diversifying selection of *H2A.P* and possibly *H2A.B*. Since the only function demonstrated for short H2A variants is in protamine deposition (Barral, et al. 2017), we speculate that the functional constraints acting on the short histone variants may be similar to those that act on protamines, which are themselves considered to be among the fastest-evolving proteins in mammalian genomes (Retief and Dixon 1993; Wyckoff, et al. 2000; Torgerson, et al. 2002; Martin-Coello, et al. 2009). Although the high dN/dS values of protamine proteins may be artificially inflated due to selective pressure to maintain high basic amino acid content across the whole protein (Rooney and Zhang 1999; Clark and Civetta 2000; Rooney, et al. 2000), these arguments cannot explain other rapidly-evolving sperm-specific DNA-packaging proteins in animal genomes (Wyckoff, et al. 2000; Kimura and Loppin 2016).

Genes encoding the short H2A variants are found almost exclusively on the X chromosome, which experiences X-inactivation and genomic imprinting (Graves 2016). This genomic location might simply be the result of evolutionary history, since that is where the genes first arose. However, the X chromosome is a curious location for genes that are exclusively expressed in the male germline, in which most X-linked genes are silenced during meiotic sex chromosome inactivation (Turner 2015). Sex chromosome linkage brings with it a suite of unconventional selective pressures that might shed light on the function of short H2A variants. Because X-linked genes spend more evolutionary time in females (XX) than males (XY) they are subjected to varying selection regimes depending on their expression bias. Yet, X-linked genes expressed in males are fully exposed to selection in the hemizygous state, and male-beneficial mutations can reach fixation even if they are harmful to females (Rice 1984; Ellegren and Parsch 2007). This mode of selection can drive the X chromosome to accumulate sex-biased genes engaged in sexual antagonism (*i.e*., beneficial to one sex and detrimental or neutral to the other). Known male-beneficial genes in placental mammals have been shown to amplify within the X chromosome and to undergo gene conversion (Ross, et al. 2005; Ellegren 2011). Interestingly, one recent mouse *H2A.L* duplicate has relocated to an autosome and two have relocated to the Y chromosome. This pattern is expected when male-beneficial genes are trying to escape female compensatory suppression (*i.e*., meiotic sex chromosome inactivation) (Ellegren and Parsch 2007).

The evolutionary signatures uncovered for *H2A.B*, *H2A.L, H2A.P* and *H2A.Q*, together with their male germline expression, suggest that they may be engaged in sexual antagonism. It is also conceivable that they participate in post-meiotic drive in spermatogenesis. Due to their X-linkage and expression during spermatogenesis, short histone variants could specifically package the chromatin of X chromosome-containing germ cells to protect against the action of a Y-linked factor. Such a mechanism has been invoked to explain the recent, dramatic acquisition of chromatin genes on both the mouse X and Y chromosomes (Bachtrog 2014; Soh, et al. 2014; Moretti, et al. 2017). Although the nature of this genetic conflict is still unclear, we hypothesize that some form of evolutionary arms race could have led to the rapid evolution of short histone H2A variants. Our evolutionary studies thus firmly establish the evolutionary novelty in the origin and diversification of short histone H2A variants, and highlight their unusual evolutionary signatures as impetus for their functional characterization.

## Acknowledgments

We are grateful to Tera Levin, Michelle Hays and Paul Talbert for comments on the manuscript, to Srinivas Ramachandran and Courtney Schroeder for help analyzing the nucleosome crystal structure, and to Mike Doud for suggesting the Jensen-Shannon distance metric. This work was supported by a postdoctoral fellowship awarded to A.M. by the Damon Runyon Cancer Research Foundation (DRG:2192-14) and grants from the NIH R01 GM074108 and from the Howard Hughes Medical Institute to H.S.M. The funders played no role in study design, data collection and interpretation, or the decision to publish this study. H.S.M. is an Investigator of the Howard Hughes Medical Institute.

## Materials and Methods

### Identification of short H2A variant genes

Canonical H2A, macroH2A, H2A.X and H2A.Z mRNA and protein sequences from representative species were retrieved from the NCBI histone database (https://www.ncbi.nlm.nih.gov/projects/HistoneDB2.0/)(Draizen, et al. 2016). In the first round of our search for short H2A sequences, we queried the assembled genomes of 25 mammals using tBLASTn (Altschul, et al. 1990; Altschul, et al. 1997). Our query sequences were human H2A.B.1.2 (NP_542451.1), rat H2A.L.2 (XP_002730244.1) and human H2A.P (NP_036406.1). Some subsequent searches used the horse H2A.L.3 sequence we identified (Supp. Table 1) as a query, because it is more conserved than rat H2A.L. Nucleotide hits were retrieved, with enough flanking sequence that the full ORF would be included, and their assembly coordinates recorded. We used the UCSC Genome Browser (Kent, et al. 2002) to identify 5’ and 3’ flanking genes for synteny analyses, or determined locations of the expected flanking genes by BLASTn. Candidate short H2A variants were further processed in two steps: 1) in silico translation to annotate putative intact ORFs and pseudogenes 2) sequence trimming after alignment to other H2A sequences manually, or using MUSCLE or MAFFT aligners (Katoh, et al. 2002; Edgar 2004).

We classified a hit as a putative short H2A if any of the following conditions were met: 1) the ORF was shorter than canonical H2As, and less than 80% identical to any other known H2A variant; 2) the ORF was found in a location that shows shared synteny with other putative short H2As; 3) the ORF shared greater similarity with other putative short H2As than any other variant. Finally, we used a combination of synteny and phylogenetic placement to classify ORFs into one of the four short H2A families: H2A.B, H2A.L, H2A.P or H2A.Q. We gave each identified short H2A variant a name that reflects the variant type (B, P, Q or L) and the ancestral syntenic locus in which it is found (1, 1.1, 1.2, 2 or 3) or with an extension reflecting the species name and an arbitrary number if found outside known syntenic loci or on short assembly scaffolds that lack flanking genes. For example, Cat *H2A.B.1.1* resides in the first H2A.B locus, and Cheetah *H2A.L.cheetah1* is on a very short genomic sequence, so flanking genes cannot be identified.

For detailed re-analysis of some clades, or of loci where syntenic H2A genes were not initially found, we used more closely-related sequences as queries for BLASTn or tBLASTn searches (*e.g*., we identified sheep and cow *H2A.P* pseudogenes using pig *H2A.P* as query in a BLASTn search, and for the more rapidly-evolving rodent clade we used mouse protein sequences as queries).

### Phylogenetic analyses

*H2A phylogeny* (FIgure 3): we used ClustalW (Larkin, et al. 2007) to align predicted protein sequences of canonical H2A, H2A.Z, H2A.X, macroH2A and short H2As. We trimmed the alignment to the histone fold domain (HFD) using published structural annotations (Luger, et al. 1997; Draizen, et al. 2016). We estimated a phylogenetic tree using maximum-likelihood methods implemented in PhyML (Guindon and Gascuel 2003; Guindon, et al. 2010) with the Jones-Taylor-Thornton substitution model (Jones, et al. 1992), 100 bootstrap replicates and optimizing tree topology, length and substitution rate.

*H2A.B, H2A.L, H2A.P and H2A.Q phylogenies*: nucleotide sequences were aligned using MUSCLE (Edgar 2004) or by hand using protein alignments as a guide. Phylogenetic trees were built by maximum-likelihood methods (PhyML) (Guindon, et al. 2010) using the HKY85 substitution model with 100 bootstrap replicates. Sequences were analyzed for evidence of recombination using the GARD algorithm implemented at datamonkey.org (Kosakovsky Pond, et al. 2006).

### Logos and nucleosome structure

Logo plots were generated using weblogo.berkley.edu (Crooks, et al. 2004) from alignments that included a single H2A.B, H2A.L, H2A.P and canonical H2A representative from each of the following species: mouse, rat, Chinese hamster, pig, panda, leopard, rhinoceros and armadillo. The isoelectric point for each of the protein was computed using the ExPASy portal (Artimo, et al. 2012) and then averaged. We displayed the published nucleosome structure (PDB:1AOI) (Luger, et al. 1997) with some residues highlighted using UCSF’s Chimera software (Pettersen, et al. 2004).

In order to obtain a quantitative view of conservation between different H2A variants, we calculated a two-way Jensen-Shannon distance metric at each amino acid position in the histone fold domain following a published formula (Doud, et al. 2015). We also adapted the calculation to allow a three-way comparison (H2A.B vs. H2A.L vs. H2A.P), calculating distance as follows:

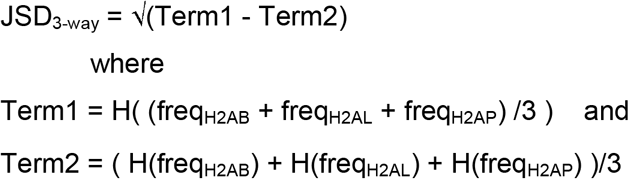

where “freq” denotes a vector of length 20 representing amino acid frequencies at each position, and H denotes the Shannon entropy of such a vector.

### Histone fold prediction of H2A.Q

We analyzed of pig and human H2A.Q sequences using HistoneDB 2.0’s “analyze your sequence” option (https://www.ncbi.nlm.nih.gov/research/HistoneDB2.0/index.fcgi/analyze/) to compare each sequence against curated HMMs representing various histone variants. We also used HHpred (Soding, et al. 2005) implemented at https://toolkit.tuebingen.mpg.de/#/ to determine whether pig and human H2A.Q sequences are likely to encode good histone folds.

### RNA-seq analysis

In order to examine expression of the short H2A variants, we analyzed publically available transcriptome data. Human, opossum and platypus data have been published (Brawand, et al. 2011), dog data are from the Broad Institute (unpublished) and pig data are from Wageningen University’s FAANG project or as published (Yang, et al. 2017). SRA identifiers are listed in Supp Table 3. We downloaded fastq files using NCBI’s SRA toolkit (https://www.ncbi.nlm.nih.gov/books/NBK158900), and mapped reads to same-species genome assemblies using tophat2 (Kim, et al. 2013), using the “–max-multihits 1” option so that multiply-mapping reads were assigned randomly to a single location. We then used bedtools multicov (Quinlan and Hall 2010) and the coordinates given in Supp Table 1 to count reads overlapping each short H2A ORF, and divided those counts by the total number of mapped reads in each sample in millions as a rough form of normalization.

### Vista plots

To search for longer stretches of similarity in the loci flanking short H2A variant genes, we used dotter (Sonnhammer and Durbin 1995) to roughly define the extent of homology, and mVISTA plots (Frazer, et al. 2004) to examine homology in more detail.

### Calculating a rate of protein divergence for canonical and variant histones

We calculated pairwise identities between H2A variants and cenH3 from alignments of full-length protein sequences to either human, dog or Tasmanian devil reference sequences. Because macroH2A contains a large added C-terminal domain, we included only the HFD for better comparison with other H2As. We obtained species divergence times from the TimeTree database (www.timetree.org), which provides estimates based on median values from numerous published studies (Hedges, et al. 2015).

### PCR amplification and sequencing of primate H2A.B and H2A.P

To obtain additional primate *H2A.B* and *H2A.P* sequences we amplified genomic DNA extracted from the following sources. Coriell biorepository: siamang gibbon, *Hylobates syndactylus* (fibroblast PR00722, and DNA KB11539); black-and-white colobus, *Colobus guereza* (fibroblast PR00980); talapoin, *Miopithecus talapoin* (fibroblast PR00716); woolly monkey, *Lagothrix lagotricha* (DNA NG05356); spider monkey, *Ateles geoffroyi* (DNA NG05352); white-faced saki, *Pithecia pithecia* (fibroblast PR00239, DNA KB5932). Frozen Zoo, San Diego Zoo: howler monkey, *Alouatta sara* DNA (OR749).

H2A.B genes from talapoin, siamang gibbon and black and white colobus were amplified (45 cycles) using PCR high fidelity Super Mix from Invitrogen following the manufacturer’s instructions, using primer sequences GCTAGGATACACAGTACTGGACGCGG (F) and GGTGGGTGGACGAGTGGACC (R).

H2A.B genes from spider monkey, howler monkey, woolly monkey and white-faced saki were amplified using nested PCR. 20 cycles using external primers were followed by 30 cycles with internal primers, using Taq DNA polymerase from Roche, following the manufacturer’s instructions. Primer sequences:

external_F–TGAGGGGCTGGGCCCTCGTCGCTAGGATACAC;

external_R-CCAAGGCAAAGTGACTGAAACAGGCAG;

internal_F-CTAGAGCAAGCCGAGCCGAGAT;

internal_R-GCTGGGGGATTTGGGGCTGGGT.

H2A.P sequences from white-faced saki and woolly monkey were amplified using Taq DNA polymerase from Roche, following the manufacturer’s instructions. Primer sequences:

P_flank_F-GAGACCTGCTCAAGCAGGAGAATCAAGC;

P_flank_R-CCAAGGCAAAGTGACTGAAACAGGCAG.

All PCR products were TOPO-TA cloned into the PCR4 vector from Invitrogen, according to the manufacturer’s instructions. Purified plasmids were Sanger-sequenced using M13 primers. Sequences are in GenBank: accessions are found in Supp. Table 1.

### Analysis of evolutionary selective pressures

We used the codeml algorithm from the PAML suite (Yang 1997) to test for positive selection on *H2A.B, H2A.Q* and *H2A.P* in simian primates, and on H2A.Q in Cetartiodactyla. Codon alignments were generated as described above, or using the online software PAL2NAL (Suyama, et al. 2006). Alignments were used as input to codeml along with either the accepted species tree (Perelman, et al. 2011), or a tree generated from the alignment using maximum-likelihood methods (PhyML) (Guindon, et al. 2010) with the HKY85 substitution model. To determine whether *H2A.B, H2A.Q* and *H2A.P* evolve under positive selection, we compared either of two “NSsites” evolutionary models that do not allow dN/dS to exceed 1 (M7 or M8a) to a model that allows dN/dS > 1 (M8). Positively selected sites were classified as those sites with M8 Bayes Empirical Bayes posterior probability > 90%. The results we present are from codeml runs using the F3x4 codon frequency model, and initial omega 0.4. Analyses were robust to use of different starting parameters (codon frequency model F61; starting omega 1.5). We also used the FUBAR program implemented at datamonkey.org (Delport, et al. 2010). One major caveat is that most of the identified codons overlap CpG dinucleotides in one or more of the aligned species. DNA methylation at cytosines within CpG sites greatly increases the rate of spontaneous deamination, causing C-to-T transition mutations (Duncan and Miller 1980; Ehrlich, et al. 1990). This increased mutation rate can artificially alter measured dN/dS ratios without the need to invoke positive or purifying selection as an explanation, and is not accounted for by currently available methods to test for positive selection. However, it remains quite possible that amino acid changes resulting from mutations at CpG sites can contribute to functional variation that selective pressures can act upon. Performing a much more conservative analysis after masking all codons overlapping CpG sites in any of the aligned species yielded no support for positive selection (not shown). However, because *H2A.B* and *H2A.P* coding sequences are GC-rich (65% and 46% respectively), masking these sites is very conservative; we removed roughly half all codons in the already-short alignments, greatly reducing the statistical power of methods to detect positive selection.

We also used codeml’s model 0 to test various alignments for signatures of purifying selection: model 0 assumes a single dN/dS value for all sites of the alignment. We estimated a phylogeny for each alignment as above, and compared the maximum likelihood of model 0 where dN/dS is freely estimated versus model 0 where dN/dS is fixed at 1 (neutral evolution).

For evolutionary analysis of the selective pressures acting on primate *H2A.L* genes (Supp. Fig. 1) we first codon-aligned primate *H2A.L* nucleotide sequences and pig *H2A.L* genes as outgroups. We used PHYML to estimate a phylogeny, specifying the HKY85 evolutionary model, 100 bootstrap replicates, and allowing PHYML to estimate equilibrium base frequencies as well as the shape of the gamma distribution (Supp. Fig. 1A). To examine selective pressures (Supp. Fig. 1B) we eliminated obvious pseudogenes from the alignment (those with stop codons or frameshifts interrupting the typical coding region), added more *H2A.L* genes from additional representative mammals, and made another phylogenetic tree. We used this alignment and tree as input into PAML’s codeml algorithm to perform branch tests of dN/dS, specifying branch model=2 (where branches of the tree are partitioned into two classes, the “foreground” and “background” branches) and NSsites=0 (so that dN/dS does not vary over sites). For each test depicted in Supp. Fig. 1B, we specified the indicated group of branches as the foreground braches, and ran codeml twice. In one run, the dN/dS values are freely estimated for both foreground and background branches. In the second run, dN/dS is fixed at 1 for the foreground branches (but is still estimated for background branches). We obtain a p-value supporting the hypothesis that the foreground branches are evolving non-neutrally by calculating twice the maximum likelihood difference between the two runs, and comparing that value to a chi-squared distribution with one degree of freedom.

## Supplementary material

### Supplementary Figure 1

Evolutionary analysis of primate *H2A.L* sequences.

Analysis of selective pressures on primate *H2A.L* sequences suggests that hominoids and Old World monkeys lack functional *H2A.L* sequences, but that New World monkeys likely retain functional H2A.L proteins. **(A)** Nucleotide phylogeny of primate *H2A.L* nucleotide sequences and pig outgroups, showing all bootstrap values >50%. The *H2A.L.2* locus acquired a stop codon in the common ancestor of all primates, and has evolved on an independent trajectory since that time. The status of the ORFs in the *H2A.L.1* and *H2A.L.3* loci is less clearcut. Although a few sequences do exhibit inactivating mutations, others appear intact, with some encoding an unusually long C-terminal extension to the protein (labeled “longORF”). **(B)** Nucleotide phylogeny of selected primate and outgroup H2A.L genes, excluding those with obvious inactivating mutations, and showing all bootstrap values >50%. We used PAML’s codeml algorithm to estimate dN/dS on selected sets of branches and to test the idea that those branches are evolving non-neutrally. While New World monkey *H2A.L* genes are somewhat likely evolve under purifying selection (dN/dS=0.62, p-value for non-neutral evolution=0.054); Old World monkey/hominoid *H2A.L* sequences evolve at a rate that cannot be distinguished from neutral evolution (dN/dS=0.79, p=0.35).

### Supplementary Figure 2

Nucleotide phylogeny of rodent *H2A.L* genes.

Evolutionary analysis shows that the three ancestral *H2A.L* loci evolve on independent trajectories in rodents. We show showing all bootstrap values >50%, with selected other lower values shown in parentheses. Many genes in the *H2A.L.3* locus are pseudogenes in rodents. The gene at the *H2A.L.2* locus has experienced additional duplications in the Mus lineage.

### Supplementary Figure 3

Alignment or Sclater’s lemur *H2A.Q* pseudogene sequence with human and pig H2A.Q genes shows that the ancestral histone fold domain had no deletions, and may suggest frameshifts in human H2A.Q.

We show the DNA sequence of the H2A.Q pseudogene in Sclater’s lemur (top), with a translation in all three frames immediately below, along with predicted protein sequences from pig and human H2A.Qs. The reading frame that aligns to intact H2A.Q genes, like pig H2A.Q, is highlighted in yellow, and putative frameshifts in the human sequence are highlighted in grey.

### Supplementary Table 1

Coordinates of all short H2A variant genes identified in this study.

### Supplementary Table 2

Detailed table of results from all analyses of selective pressures.

### Supplementary Table 3

List of SRA datasets used for RNA-seq analysis.

### Supplementary Table 4

Data used for Figure 8A.

**Supplementary Data 1: Alignment of histone fold domains used to generate the phylogeny shown in Figure 3.**

**Supplementary Data 2: Alignments used to generate logo plots shown in Figure 4.**

**Supplementary Data 3: Alignments used for selection analyses.**

**Supplementary Data 4: Alignments used for gene conversion analyses shown in Figure 5.**

